# Improved Production of Taxol® Precursors in *S. cerevisiae* using Combinatorial *in silico* Design and Metabolic Engineering

**DOI:** 10.1101/2023.06.11.544475

**Authors:** Koray Malcı, Rodrigo Santibáñez, Nestor Jonguitud-Borrego, Jorge H. Santoyo-Garcia, Eduard J. Kherkoven, Leonardo Rios-Solis

**Affiliations:** Institute for Bioengineering, School of Engineering, University of Edinburgh, King’s Buildings, Edinburgh EH9 3BF, United Kingdom; Centre for Engineering Biology, University of Edinburgh, King’s Buildings, Edinburgh EH9 3BF, United Kingdom; Department of Pediatrics, University of California, San Diego, La Jolla, CA, 92093-0760, USA; Department of Life Sciences, Chalmers University of Technology, Kemivägen 10, SE412 96, Gothenburg, Sweden; School of Natural and Environmental Sciences, Molecular Biology and Biotechnology Division, Newcastle University, Newcastle upon Tyne NE1 7RU, United Kingdom; Department of Biochemical Engineering, The Advanced Centre for Biochemical Engineering, University College London, Gower Street, London, WC1E 6BT, UK

**Keywords:** *in silico* design, computational metabolic engineering, yeast model, synthetic biology, Taxol production

## Abstract

Integrated metabolic engineering approaches combining system and synthetic biology tools allow the efficient designing of microbial cell factories to synthesize high-value products. In the present study, *in silico* design algorithms were used on the latest yeast genome-scale model 8.5.0 to predict potential genomic modifications that could enhance the production of early-step Taxol® in previously engineered *Saccharomyces cerevisiae* cells. The solution set containing genomic modification candidates was narrowed down by employing the COnstraints Based Reconstruction and Analysis (COBRA) methods. 17 genomic modifications consisting of nine gene deletions and eight gene overexpression were screened using wet-lab studies to determine whether these modifications can increase the production yield of taxadiene, the first metabolite in the Taxol® through the mevalonate pathway. Depending on the cultivation condition, most of the single genomic modifications resulted in higher taxadiene production. The best-performing strain, named KM32, contained four overexpressed genes, *ILV2, TRR1, ADE13* and *ECM31*, from the branched-chain amino acid biosynthesis, thioredoxin system, *de novo* purine synthesis, and the pantothenate pathway, respectively. Using KM32, taxadiene production was increased by 50%, reaching 215 mg/L of taxadiene. The engineered strain also produced 43.65 mg/L of taxa-4(20),11-dien-5α-ol (T5α-ol), and 26.2 mg/L of taxa-4(20),11-dien-5-α-yl acetate (T5αAc) which are the highest productions of these early-step Taxol® metabolites reported until now in *S. cerevisiae*. The findings of this study highlight that the use of computational and integrated approaches can ensure determining promising modifications that are difficult to estimate intuitively to develop yeast cell factories.

## INTRODUCTION

Microbial chassis have been intensely studied to produce industrially essential compounds such as pharmaceuticals [1], biofuels [2], enzymes [3] and polymers [4] in sustainable and economically attractive ways. Among these organisms, baker’s yeast, *Saccharomyces cerevisiae,* is the most widely studied eukaryotic synthetic biology chassis [5]. In relatively cheap media, *S. cerevisiae* can produce high biomass [6], and its fermentation can be efficiently controlled and scaled up [7]. Also, many synthetic biology tools have been developed, and genetic parts have been characterized to design yeast strains in a shorter time and in a more reliable way [8, 9].

*S. cerevisiae* has been used as a platform to produce high-value biopharmaceuticals, ranging from recombinant therapeutic proteins to plant-derived natural products [10, 11]. By integrating heterologous plant-derived genes, early-step precursors of Taxol® (paclitaxel), a leading anticancer drug with a market size over billions of USD [12], have been produced by yeast cell factories [13–17]. Geranylgeranyl diphosphate (GGPP), a product of the yeast mevalonate pathway, is the substrate of the cyclisation step, the first step in the Taxol® biosynthesis pathway catalyzed by the taxadiene synthase [18]. Once GGPP is converted into taxadiene (taxa-4(5),11(12)-diene), it is followed by a hydroxylation step catalyzed by a class II cytochrome P450 hydroxylase, taxadiene-5α-hydroxylase (T5αOH) with the help of cytochrome P450 reductase (CPR), and an acylation step catalyzed by taxadiene-5α-ol-O-acetyltransferase (TAT), respectively [12]. Although these enzymes have been expressed, and their products have been successfully produced in yeast [19, 20], there is still room to improve their titers for economically feasible production of the subsequent Taxol® precursors. Figure 1 illustrates the biochemical reactions in the mevalonate and Taxol® pathways.

**Figure 1:**
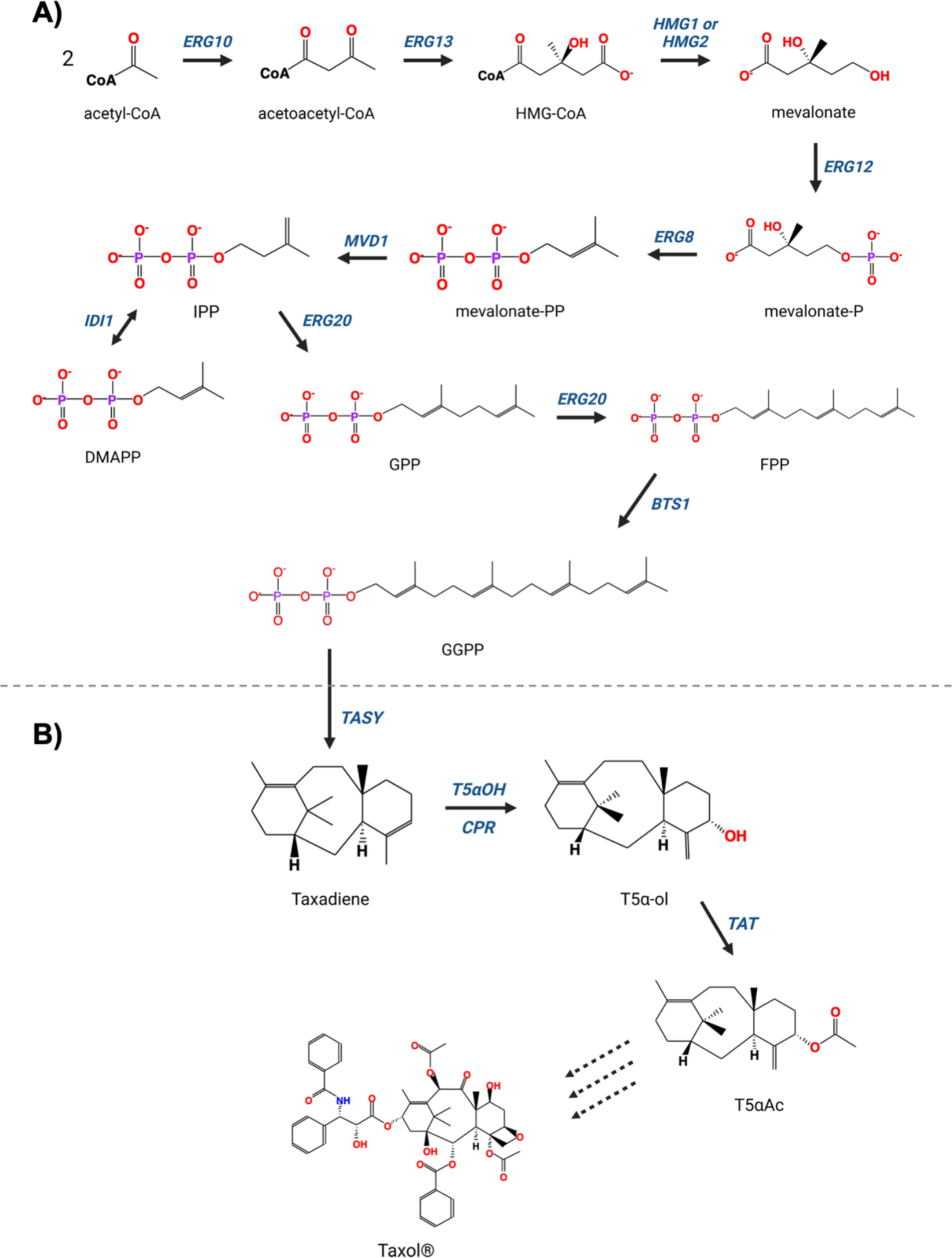
**A)** The native biochemical reactions in the mevalonate pathway leading to the production of GGPP in yeast. **B) The** Taxol® biosynthesis from GGPP in yew trees. HMG-CoA: hydroxymethylglutaryl-CoA, mevalonate-P: R-5-hosphomevalonate, mevalonate-PP: R-5 diphosphomevalonate, IPP: isopentenyl diphosphate, DMAPP: dimethylallyl diphosphate, GPP: geranyl diphosphate, FPP: farnesyl diphosphate, GGPP: geranylgeranyl diphosphate, T5α-ol: taxa-4(20),11-dien-5α-ol, T5αAc: taxa-4(20),11-dien-5-α-yl acetate. The dashed arrows represent naturally occurring multiple steps.

As omics technologies are evolving and the sequencing platforms are becoming more accessible, genome-scale metabolic models (GEMs) are being reconstructed for many organisms to convert the integrated omics data into valuable system-level information [21]. Since the first yeast GEM was reconstructed in 2003, many alternative and improved versions of *S. cerevisiae* GEMs have been developed by adding and connecting more genes, reactions, and compartments [22–24]. The latest consensus yeast GEM, yeast 8 [24], is currently being used to construct more up-to-date and corrected versions of yeast GEMs to mimic the yeast metabolism better. Among the yeast 8 derivatives, yeast 8.5.0 includes 4055 reactions, 2742 metabolites, and 1151 genes in 14 cellular compartments [25]. As a bottom-up system biology tool, yeast GEMs have been employed in many applications, from designing yeast cell factories to optimizing culture conditions [26]. In recent years, the coupling of GEMs and constraint-based reconstruction and analysis (COBRA) methods [27] has found widespread usage in biological applications. COBRA is an integrative analysis framework that can be applied to biochemical systems to demonstrate and predict the connections from phenotype to genotype by imposing constraints considering factors such as physicochemical laws, environmental conditions, and genetic information [28, 29].

We previously reported the production of early-step Taxol® precursors through the improved mevalonate pathway using our engineered yeast strains [19, 20]. In the present study, we used a combinatorial *in silico* system biology approach to further improve the flux towards GGPP in the mevalonate pathway to enhance the titers of the taxadiene and the next precursors in the Taxol® pathway. We applied three strain design frameworks, OptKnock [30], OptGene [31] and OptForce [32] on the latest yeast GEM, yeast 8.5.0, to predict gene and reaction candidates to be knocked out or overexpressed. The genomic modifications determined by *in silico* analyses were implemented using the CRISPR-based ACtivE toolkit [33] to design yeast strains. In total, the knocking-out of nine genes and the overexpression of eight genes, as well as multiple combinations were tested. Following the screening of the strains, the best-performing one was also used in 250 mL in a mini-scale bench-top bioreactor. We achieved a 50% increase in taxadiene yield compared to the parent strain when four reactions were upregulated by integrating an extra copy of *ILV2, TRR1, ADE13* and *ECM31* genes driven by a galactose-inducible promoter. Utilizing a micro-scale high-throughput bioreactor platform, BioLector, we detected 215 mg/L of taxadiene, 35.4 mg/L of T5α-ol and 26.2 mg/L of T5αAc in complete synthetic media (CSM). These are the highest titers reported until now in *S. cerevisiae*. Our findings showed that the genomic modifications used in this study can be applied to improve the metabolic flux towards the yeast mevalonate pathway to produce high-value terpenoids.

## MATERIALS & METHODS

### *in silico* Design and Analyses

Three *in silico* design methods, OptKnock, OptGene and OptForce, were used to find the potential genomic modifications to increase the concentration of Taxol® precursors. Cytosolic acetyl-CoA and geranylgeranyl diphosphate (GGPP) were separately selected as target metabolites to increase production in glucose or galactose-containing media. To mimic CSM, uptake routes for key molecules such as inorganic phosphate, sulphate, ammonia, and oxygen were unconstrained, while secretion routes for acetate, carbon dioxide, ethanol, glycolaldehyde, diphosphate, water, and glycerol and acetaldehyde were enabled. When galactose exchange was enabled, glucose exchange was constrained or vice versa.

After the design algorithms suggested genomic modification candidates, several analysis methods were used to predict high-potential modifications. Flux variability analysis (FVA) [34] was used to find the maximum flux differences towards the target metabolite between wild-type and mutant strains. Metabolite interaction networks were constructed [35, 36], and fluxes differences were checked on the metabolites in the mevalonate pathway.

The COBRA Toolbox v3.0 [29, 37] was used in MATLAB R2019a for *in silico* studies. Gurobi Optimizer (9.5.0) was used as the solver in MATLAB R2019a. Yeast GEM 8.5.0 [25] was used for yeast metabolism. The ECDF Linux Compute Cluster (Eddie), the University of Edinburgh’s research computing cluster, was employed to find to run OptForce. Escher [38] was used on the iMM904 yeast model to visualize the metabolic maps since the yeast 8 does not have a compatible file format with Escher.

### Oligonucleotides and Reagents

All primers used in the study are listed in Table S1 and S3. The primers were ordered from Integrated DNA Technologies (IDT) as standard DNA oligos. Synthetic gRNA cassettes (Table S2) were ordered from Twist Bioscience. Phusion Flash High-Fidelity PCR Master Mix (Thermo Scientific™) and PrimeSTAR® GXL DNA Polymerase (TaKaRa) were used for PCR reactions to produce the DNA parts for genome modifications. GeneJET PCR Purification Kit (Thermo Scientific™) was used for PCR clean-up. DreamTaq Green PCR Master Mix (Thermo Scientific™) was used for colony PCR.

### Strains and Media

The original *S. cerevisiae* strain used in this study is a CEN.PK2-1C-originated yeast strain, *LRS6* (*MATa, leu2-3, 112::HIS3MX6-GAL1p-ERG19/GAL10p-ERG8; ura3-52::URA3-GAL1p-MvaS^A110G^/GAL10p-MvaE*; *his3Δ1::hphMX4-GAL1p-ERG12/GAL10p-IDI1; trp1-289::TRP1_GAL1p-CrtE (X. dendrorhous)/GAL10p ERG20; YPRCdelta15::NatMX-GAL1p-CrtE/GAL10p-CrtE*; *ARS1014::GAL1p-TASY-GFP*; *ARS1622b::GAL1p-MBP-TASY-ERG20*; *ARS1114a::TDH3p-MBP-TASY-ERG20; ARS511b::GAL1p-T5αOH/GAL3-CPR; RKC3::GAL1-TAT*) [20]. Briefly, the original strain contains additional copies of native and heterologous genes in the mevalonate pathway to improve GGPP production, along with codon-optimized heterologous genes encoding the enzymes responsible for early steps in Taxol® biosynthesis (Figure 1 and Figure 3A). An additional *GAL1* promoter-driven *TAT* gene was integrated to the *RKC4* location into the *LRS6* genome, which was used as the parent strain in the subsequent studies. The other strains derived from the parent strain are shown in Table S4.

**Figure 2:**
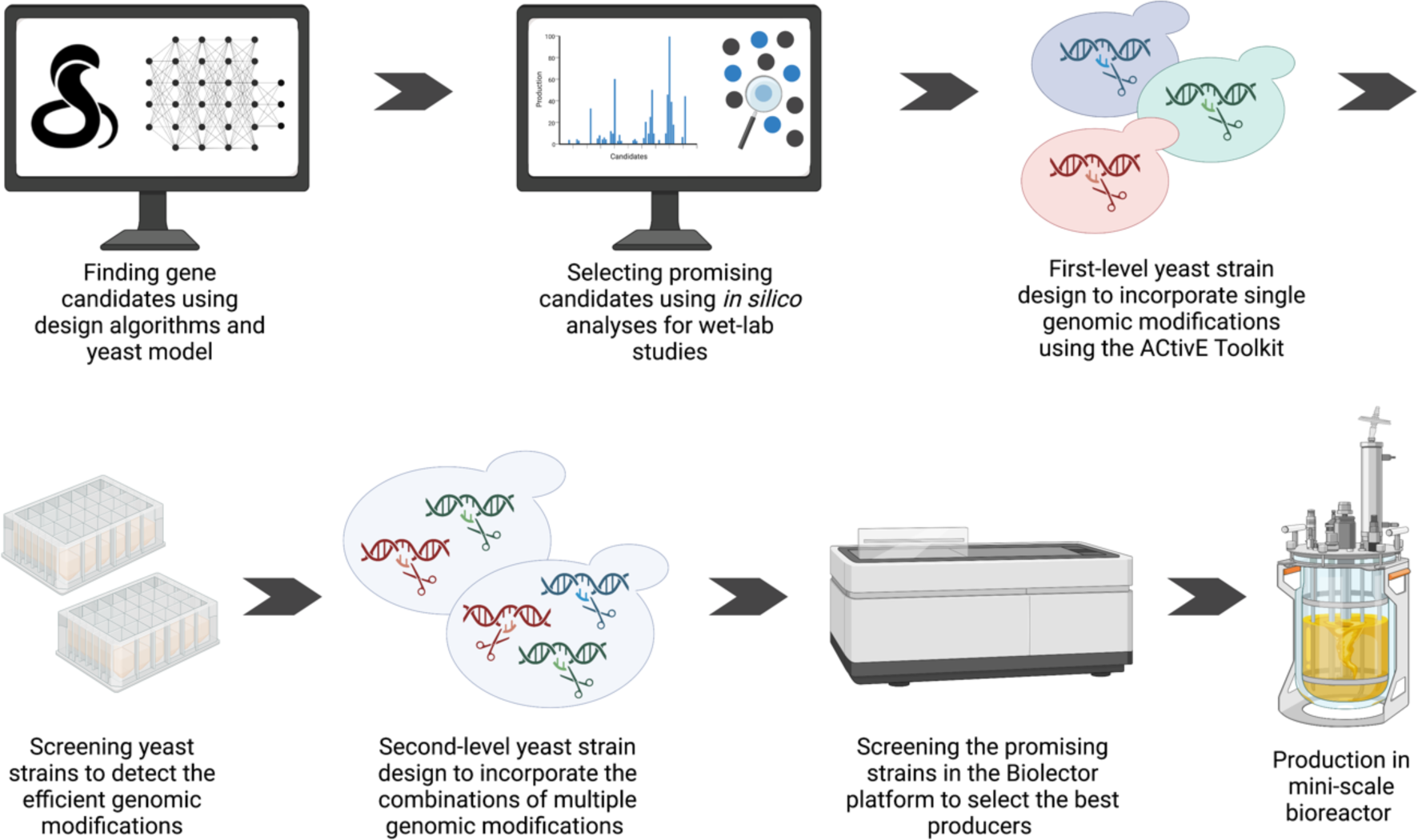
The workflow followed in this study from *in silico* design to lab bench scale production. Three design algorithms, OptKnock, OptGene and OptForce, were used in the COBRA Toolbox employing the Yeast 8.5.0 GEM. After selecting the gene candidates, the yeast strains containing single genomic modifications were designed. To engineer the yeast genome, the ACtivE Toolkit was used. The strains were then screened in deep-well plates to detect the good performance genomic modifications. These modifications were then combined to design second-level yeast strains containing multiple genomic modifications. The best-performing strains were then screened in the Biolector platform with real-time pH, DO and biomass monitoring. Finally, the best-producer was used for the productions a lab bench scale bioreactor.

**Figure 3:**
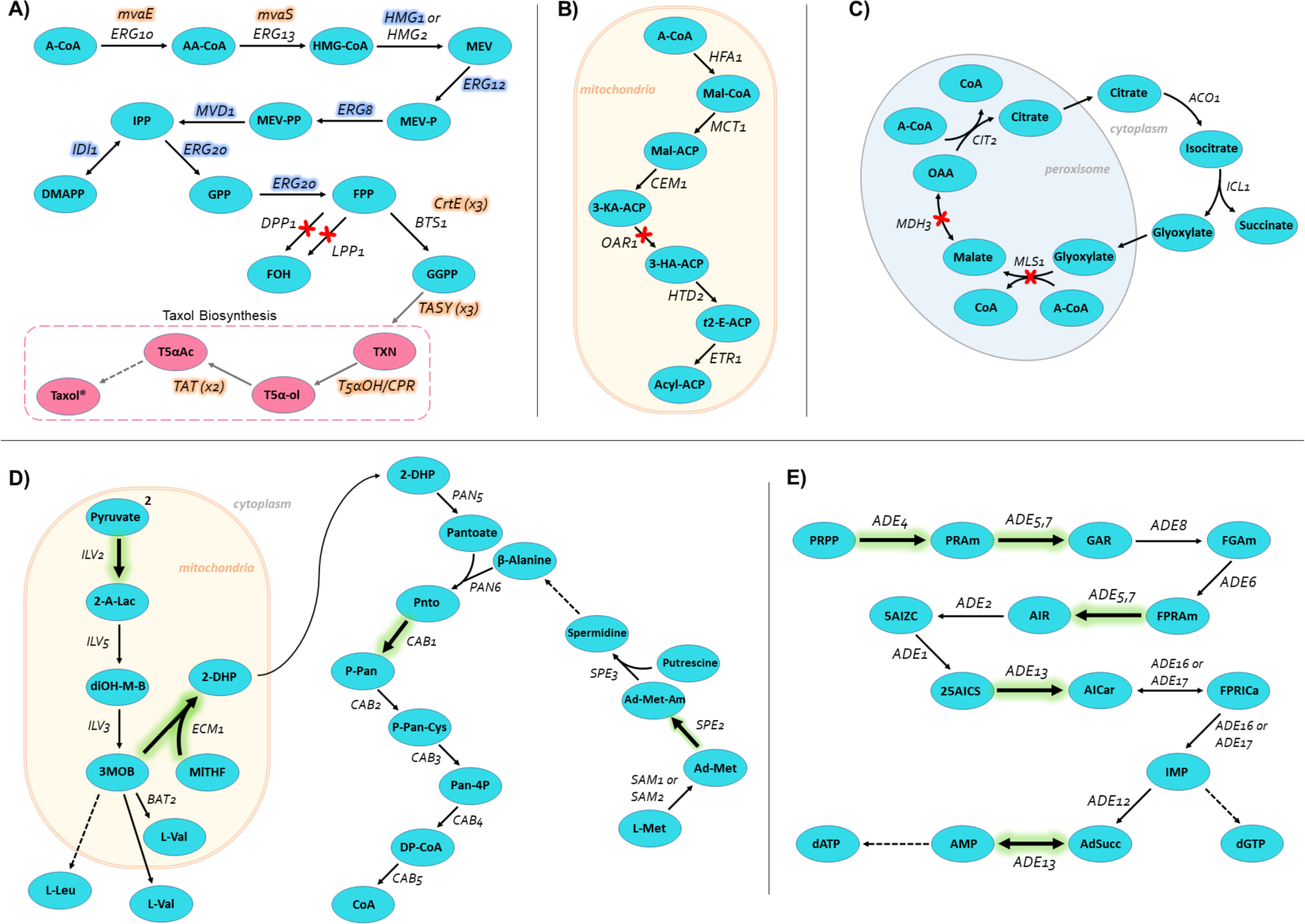
The pathways, genes and reactions modified in this study. **A)** The mevalonate pathway. The additional copies of native genes (blue shaded) and the heterologous genes (orange shaded) integrated in a previous study to construct the parent strain LRS6 [20] are highlighted. **B)** Fatty acid biosynthesis in mitochondria **C)** Glyoxylate cycle **D)** Connected pathways; branched-chain amino acid biosynthesis (from two pyruvate molecules to L-valine or L-leucine), pantothenate pathway (from 3-methyl-2-oxobutanoate a 5,10-methylenetetrahydrofolate to pantothenate) and the spermidine biosynthesis (from *S*-adenosyl-L-methionine to spermidine) **E)** *de novo* purine biosynthesis. The red crosses indicate corresponding gene deletions suggested by the design algorithms; thick arrows highlighted with green colour indicate the corresponding overexpressed genes suggested by the design algorithms. The arrows show the direction of the fluxes. Dashed arrows indicate multiple reactions. A-CoA: acetyl-CoA, AA-CoA: acetoacetyl-CoA, HMG-CoA: hydroxymethylglutaryl-CoA, MEV: mevalonate, MEV-P: R-5 phosphomevalonate, MEV-PP: R-5-diphosphomevalonate, IPP: isopentenyl diphosphate, DMAPP: dimethylallyl diphosphate, GPP: geranyl diphosphate, FPP: farnesyl diphosphate, FOH: farnesol, GGPP: geranylgeranyl diphosphate, TXN: taxadiene, T5α-ol: taxa-4(20),11-dien-5α-ol, T5αAc: taxa-4(20),11-dien-5-α-yl acetate, Mal-CoA: malonyl-CoA, Mal-ACP: malonyl-ACP, 3-KA-ACP: 3-ketoacyl-ACP, 3-hydroxyacyl-ACP, *t*2-E-ACP: *trans-*2-enoyl-ACP, OAA: oxaloacetic acid, 2-A-Lac: 2-acetolactate, diOH-M-B: 2,3-dihydroxy-3 methylbutanoate, 3MOB: 3-methyl-2-oxobutanoate, 2-DHP: 2-dehydropantoate, Pnto: R-pantothenate, P-Pan: R-4’ phosphopantothenate, P-Pan-Cys: R-4’ phosphopantothenoyl-L-cysteine, Pan-4P: 4’-phosphopantetheine, DP-CoA: 3’-dephospho-CoA, PRPP: 5-Phospho-alpha-D-ribose 1-diphosphate, PRAm: 5-Phospho-beta-D-ribosylamine, GAR: N1-(5-Phospho-D-ribosyl) glycinamide, FGAm: N2-Formyl-N1-(5-phospho-D-ribosyl) glycinamide, FPRAm: 2-(Formamido)-N1-(5-phospho-D-ribosyl) acetamidine, AIR: 5-amino-1-(5-phospho-D-ribosyl)imidazole, 5AIZC: 5-amino-1-(5-phospho-D-ribosyl) imidazole-4-carboxylate, 25AICS: (S)-2-[5-Amino-1-(5-phospho-D-ribosyl) imidazole-4-carboxamido]succinate, AICar: 5-Amino-1-(5-Phospho-D-ribosyl)imidazole-4-carboxamide, FPRICa: 5-Formamido-1-(5-phospho-D-ribosyl)imidazole-4-carboxamide

Complete synthetic medium (CSM) consisting of 0.67% (w/v) yeast nitrogen base without amino acids (Alfa Aesar™), 0.08% (w/v) complete supplement mixture (MP Biomedicals™) and 2% (w/v) glucose (Alfa Aesar™) or 2% (w/v) galactose (ACROS Organics™) was used for the cultivations. Pre-cultures were grown in standard rich medium, YPD medium, 1% (w/v) yeast extract (Fisher BioReagents™), 2% (w/v) peptone (Merck, Millipore®), 2% (w/v) glucose. To select the positive transformants, selective media, CSM-URA, 0.67% (w/v) yeast nitrogen base without amino acids, 0.077% (w/v) complete supplement mixture without uracil, 2% glucose, 2% agar (ACROS Organics™) was used.

### Yeast Transformation and Strain Construction

The chemicals and reagents were obtained from Sigma-Aldrich unless otherwise stated. Transformations were carried out according to LiAc/PEG heat-shock method [39]. Briefly, fresh cultures were prepared to obtain the cells in the exponential phase following an overnight culture. The cells were then washed using sterile water and were pelleted by centrifugation. The transformation mix, 240 μL PEG (50%(w/v)), 36 μL 1.0 M lithium acetate (LiAc) and 50 μL single-stranded carrier DNA (2.0 mg/mL, herring sperm DNA, Promega), was added onto the cell pellet. Following this, DNA fragments and water were added until the volume was made up to 360 μL. After homogenous transformation mixes were obtained, the cells were incubated for 45 minutes at 42 °C. Finally, the transformation mix was removed, the cells were plated onto the selective media, CSM-URA, and the plates were incubated for 2-3 days at 30°C.

Genomic modifications, gene integrations or deletions, were carried out using the modular ACtivE toolkit and method that was recently developed by our group [33]. 50 fmol equivalent molarity of each plasmid module was used to assemble a single all-in-one CRISPR plasmid. Four plasmid modules, Cas9 cassette, selection marker (*URA3*), storage part (bacteria ORI and *Amp^R^*) and yeast origin of replication (2μ) were combined with the corresponding gRNA cassettes according to the ACtivE method [33]. ARS209, ARS306, ARS727, ARS1531 and ARS1603 were used as integration sites for the genomic integrations [33]. 500 ng – 1000 ng from each donor DNA part containing overlapping fragments with their neighbor parts was used. The upstream homology arm (UHA), promoter, coding sequence (with terminator) and downstream homology arm (DHA) were designed for integrations, whereas only UHA and DHA were used for deletions. The potential crRNA sequences on each region were scored using CRISPOR [40] (http://crispor.tefor.net), an online gRNA selection tool giving sequence-based scores using sequence prediction algorithms.

The colonies were first screened using colony PCR to detect the genomic modifications. Genomic DNAs of the positive transformants were extracted using Pierce Yeast DNA Extraction Reagent Kit (Thermo Scientific™). The target regions were confirmed by Sanger sequencing performed at GENEWIZ, Inc (Leipzig, Germany).

Before the cultivations, *URA3* containing CRISPR plasmids were removed using the 5-Fluoroorotic Acid (5-FOA) (Thermo Fisher Scientific) counter-selection method with a synthetic defined medium supplemented with 0.1% (w/v) 5-FOA.

### High-throughput Strain Screening

To determine the best producers of the Taxol® precursors, the strains containing single genomic modifications (integration or deletion) were screened using V-shaped, 24-deep well-plates (Axygen™). A 20% dodecane (ACROS Organics™) overlay was added to set a working volume of 2 ml in each well, and CSM with glucose or galactose was used for the cultivations. The initial OD_600_ was set to 1.0 for each well by diluting pre-cultures. The plates were incubated with shaking at 350 rpm at 30 °C for 72 hours on thermomixers. Gas permeable adhesive plate seals (Thermo Scientific™) ensured oxygen transfer. The total biomass in each well was measured at 600 nm wavelength using the Nanodrop™ 2000c spectrophotometer (Thermo Scientific™).

In the second-level screening, the best performing strains with single genomic modifications and the strains containing multiple modifications were screened employing a BioLector Pro (mp2-labs) microbioreactor-screening platform. A flower-shaped, transparent bottom 48 well-plate (mp2-labs) containing pH and dissolved oxygen (DO) optodotes was used for online monitoring of biomass, pH and DO. Similar to the first screening, glucose or galactose-containing CSM was used as the medium, the initial OD_600_ was set to 1.0, and a 20% dodecane overlay was used in 1 mL working volume. The plate was covered using a gas-permeable sealing foil with an evaporation reduction layer (mp2-labs), and the temperature was maintained at 30°C under the agitation of 1000 rpm. Biomass absorbance units were measured with the gain=6.

After cultivations in the plates, dodecane layers were collected following a centrifuge step, and the taxane production was analyzed via gas chromatography-mass spectrometry (GC-MS).

### Bioreactor Cultivation

MiniBio500 bioreactor (Applikon Biotechnology) was used for larger scale cultivation for the best producer strain. 250 mL total reaction volume containing a 20% dodecane layer was used for the cultivations. Similar to the other cultivations, the initial OD_600_ was adjusted to 1.0 by diluting overnight cultures. The same medium compositions as the previous experiments, glucose or galactose containing CSM, were used. To mitigate the foam formation, polypropylene glycol P2000 0.01% (v/v) (Alfa Aesar), was used in the medium as anti-foam. pH, DO, and temperature were measured online using the my-control system (Applikon Biotechnology). Temperature was set to 30 °C. A setpoint 70% saturation of dissolved oxygen (DO) was applied, an air sparger was used automatically to maintain O_2_ level. pH was maintained in a particular range (5.0<pH<6.5) and 1 M of NaOH was added when the pH was below the threshold. Biomass was measured online using the Optura system with a BE2100 OD sensor (BugLab). Samples were taken daily for offline biomass measurement and quantification of the metabolite concentrations.

An inverted microscope (Leica Microsystems) was used following all cultures in this study to detect if there was microbial contamination. The cells were monitored under 100X lens with immersion oil (Nikon).

### Metabolite Identification and Quantification

The dodecane layer collected at the end of the cultivations was analyzed by GC–MS as described previously [19]. A GC system, Trace™ 1300 Series (Thermo Scientific™), equipped with TraceGOLD™ TG-SQC GC column, was used. The mass spectra of 50–650 m/z were recorded on an ISQ™ Series Single Quadrupole MS (Thermo Scientific™) using EI ionization mode and a scan time of 0.204 s. The GC-MS data were processed using the Xcalibur™ software. Pure taxadiene provided by the Baran Lab (The Scripps Research Institute) and GGOH (Sigma Aldrich) were used as standards. The concentrations of the additional compounds were calculated relative to the taxadiene standard.

### Statistical analysis

Strain screening experiments were conducted in at least three replicates. The well layouts in both 24-well plates and 48 well-plate were randomized to mitigate the plate effects and possible errors. The error bars represent the standard deviations of different experiments. The one-way analysis of variance (ANOVA) was used to determine if there was a statistically significant difference between the samples or experiments. The null hypothesis considered no significant difference between the samples/runs; the null hypothesis was rejected if the p-value ≤ 0.05.

## RESULTS & DISCUSSION

The main objective of the study was to improve metabolic fluxes towards the mevalonate pathway, thus, towards GGPP which is a common precursor of diterpenes [41], to enhance the production of the early step Taxol® precursors synthesized by our engineered yeast strains. To this end, computer-aided design was used to predict/identify potential genomic modifications that would be difficult to determine intuitively. Strain design algorithms were used on yeast GEM 8.5.0. The gene deletion or integration candidates were simulated using COBRA tools. The strains containing single genomic modifications were then designed using the ACtivE toolkit and were screened in V-shaped deep-well plates. The promising single genomic modifications and their combinations were tested in a BioLector microbioreactor system. Following this, the best producer was also tested in a lab bench bioreactor. Figure 2 demonstrates the whole process explained here.

### in silico Design

Our combinatorial approach benefited from three design algorithms, OptKnock, OptGene and OptForce on yeast GEM 8.5.0. We previously engineered a yeast strain, LRS6, with the modified mevalonate pathway and an increased flux towards GGPP as shown in Figure 1 and Figure 3A [20]. Here, acetyl-CoA, and GGPP, the first and last metabolites in the mevalonate pathway, respectively (Figure 3A), were targeted individually by adding corresponding exchange reactions. Since the integrated genes in LRS6 were driven by inducible galactose promoters, the simulations were carried out using CSMs that contain galactose or glucose, separately. The conditions in a standard complete synthetic medium (CSM) were mimicked, and uptake and secrete routes were constrained or enabled accordingly. Nonessential reactions and genes were first determined for OptKnock and OptGene using flux balance analysis (FBA) [42]. Minimum growth rate waw set to 50% of wild-type growth when targeting acetyl-CoA for OptKnock deletions, however, it was set to and 20% for GGPP as OptKnock could not predict potential gene deletions in favour of GGPP maintaining higher growth rate. OptGene was run over 500 and 1000 generations, and all predictions were considered. The genomic modifications suggested by both the first “must set” and the second “must set” of OptForce runs [32] were considered. Since the design algorithms, OptKnock, OptGene and OptForce, used in this study prioritise growth-coupled solutions, all solutions taken into account were growth-coupled. Table 1 compiles the selected target genes and corresponding design algorithms.

**Table 1:** Predicted genes by different design algorithms in various media conditions.

| Target Gene | Design Algorithm | Target Compound | Carbon Source | Intervention |
| --- | --- | --- | --- | --- |
| *DPP1* | OptGene | Acetyl-CoA | Glucose | Knock-out |
| *OAR1* | OptGene | GGPP | Galactose | Knock-out |
| *MDH3* | OptKnock & OptForce | Acetyl-CoA | Glucose | Knock-out |
| *ACH1* | OptKnock & OptGene | Acetyl-CoA | Glucose | Knock-out |
| *MLS1* | OptKnock | Acetyl-CoA | Galactose | Knock-out |
| *DIT1* | OptGene | Acetyl-CoA | Glucose | Knock-out |
| *LPP1* | OptGene | GGPP | Glucose & Galactose | Knock-out |
| *ISC1* | OptGene | Acetyl-CoA | Galactose | Knock-out |
| *DTR1* | OptGene | GGPP | Glucose | Knock-out |
| *ILV2* | OptForce | Acetyl-CoA | Galactose | Overexpression |
| *TRR1* | OptForce | Acetyl-CoA | Galactose | Overexpression |
| *ADE4* | OptForce | Acetyl-CoA | Glucose & Galactose | Overexpression |
| *ADE5,7* | OptForce | Acetyl-CoA | Glucose & Galactose | Overexpression |
| *ADE13* | OptForce | Acetyl-CoA | Glucose & Galactose | Overexpression |
| *ECM31* | OptForce | Acetyl-CoA | Glucose & Galactose | Overexpression |
| *CAB1* | OptForce | Acetyl-CoA | Glucose & Galactose | Overexpression |
| *SPE2* | OptForce | Acetyl-CoA | Glucose & Galactose | Overexpression |

Using different conditions and frameworks as highlighted above, several gene and reaction candidates were found for either acetyl-CoA or GGPP overproduction (Figure 3). To narrow down the possible genomic modifications for a more feasible process for strain design, first, FVA was employed to predict the effects of the suggested genomic modifications on acetyl-CoA and GGPP productions as well as the biomass. Since the model’s objective function, biomass reaction, was targeted in a standard optimization process, maximum flux values might be in favor of the biomass. Therefore, the objective functions were first removed to minimize the bias. The lower value was first set to 0.01 for the biomass function. The initial fluxes of the candidate reactions were set to 0.01 in the initial model, and the fluxes were changed to 0.00 and 0.02 in the final model to mimic deletion (reaction knock-out) and overexpression (gene integration), respectively. The genes that could not show an increase or showed a decrease for the maximum values of the objective functions (acetyl-CoA, GGPP and biomass reactions) were not considered for the further studies. Figure 4 shows the flux rates between the final and initial models of the selected genes.

**Figure 4:**
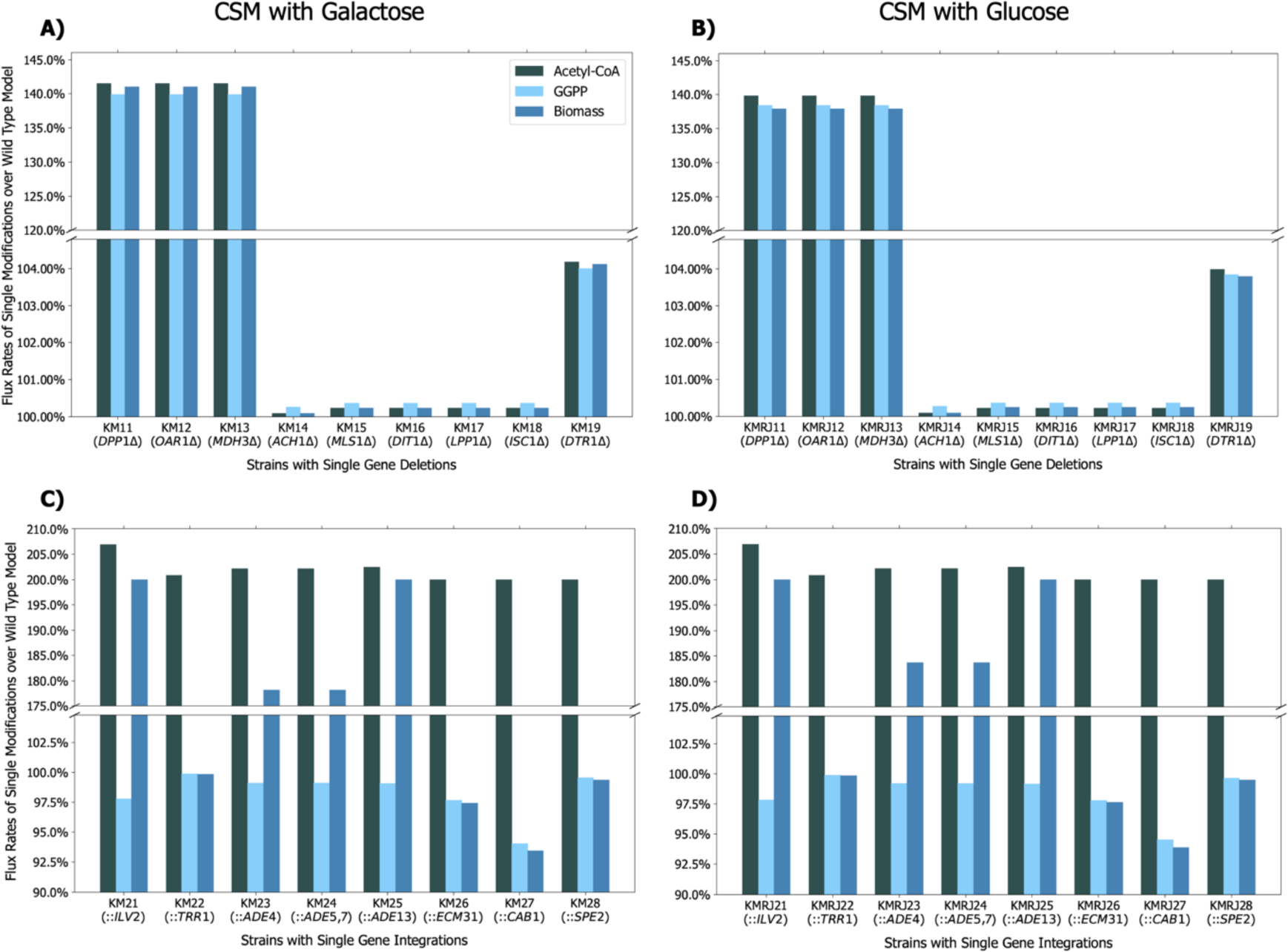
Comparison of maximum *in silico* fluxes calculated by FVA for corresponding objectives, acetyl-CoA, GGPP and biomass reactions. For simplicity, the initial state (flux value was 0.01) was referred to as wild-type, while the final state (flux value is 0.00 for deletion and 0.02 for integration) was referred to as modification. A, C were simulated in galactose containing CSM, B, D were simulated in glucose-containing CSM. Δ; gene deletion, ::; gene integration.

Although clear increases were found with approximately two-fold overproduction in acetyl-CoA when particular genes were overexpressed *in silico,* the overproduction rates were relatively low in gene deletions except for *DPP1, OAR1* and *MDH3* that showed ∼40% increase (Figure 4). To understand the lower increases better, metabolite interaction maps were constructed for the gene deletions including *ACH1, MLS1, DIT1, LPP1, ISC1* and *DTR1* (Figure 4A & 4B) to investigate the effects of these gene knock-outs. First, the wild-type model’s metabolite interactions in the mevalonate pathway were determined with their fluxes, as shown in Figure 5. The acetyl-CoA-centered and GGPP-centered interactions were investigated from this map, and flux differences between the wild-type and knock-out models were compared (Figure 5). We could not detect flux increase towards GGPP in any of the knock-out models even though the objective functions were removed, as explained above. Therefore, only acetyl-CoA increases were considered. The gene deletions that were predicted to increase fluxes in FVA (Figure 4A & 4B) and in metabolite interaction maps (Figure 5, Figure S1-S7) were selected for the strain construction.

**Figure 5:**
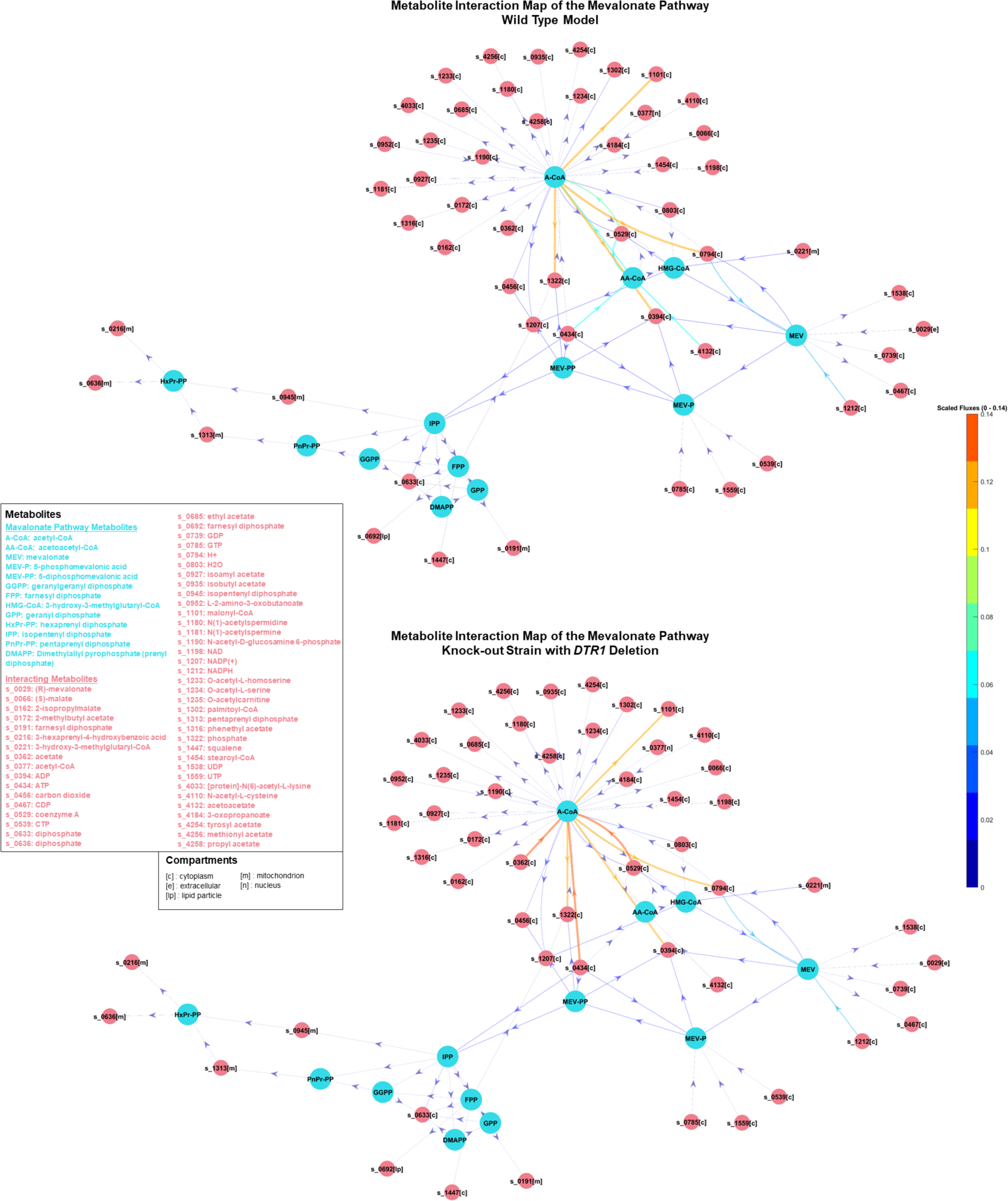
Metabolite interactions with corresponding fluxes in the mevalonate pathway in the wild-type model and the *DTR1* deleted knock-out model. Metabolite interactions were constructed using metabolite-metabolite interaction network in COBRA Toolbox [35, 36]

For overexpressions, only the reactions that increased the fluxes towards acetyl-CoA production were considered (Table 1). This was because OptForce suggested to upregulate particular reactions in the mevalonate pathway (Table S5) for GGPP overproduction, and these were already improved in our engineered strain, LRS6. OptForce did not find upregulation of early step reactions (the reactions of *ERG10* and *ERG13* genes) useful in the mevalonate pathway; the maximum GGPP production did not change in FVA compared to the initial state. It was a consistent result found by *in silico* simulations as a previous experimental study reported that early step mevalonate pathway genes of *S. cerevisiae* did not show efficient production of mevalonate compared to bacterial-sourced equivalents [43]. OptForce did not suggest any reaction in other pathways to be upregulated to enhance GGPP production.

### Strain Construction and Screening of the Single Genomic Modifications

As our engineered yeast strains contained additional genes (Figure 3) driven by galactose-inducible promoters, the parent strain KM1 (Table S4), was not able to induce the integrated mevalonate pathway genes using glucose as sole carbon source. Therefore, we first deleted the *GAL80* gene yielding with EJ1 strain (Table S4) to allow glucose utilization as Gal80 protein inhibits the transcription of the galactose-inducible genes in the absence of galactose [44]. These two strains were then designed to incorporate the genomic modifications, single-gene deletion or integration (overexpression), predicted by the design algorithms. When the additional copies of native yeast genes were integrated for overexpression, native promoters were simply changed with GAL1 promoters using the ACtivE method [33] to enhance their expression rates [19, 45]. Additional copies were integrated into an intergenic region close to the autonomously replication sequence (ARS) 1603 [33] in each strain to screen the effect of single gene overexpression.

In this study, nine gene knock-outs and eight gene overexpressions were tested to determine whether these modifications affect biomass and the production of early steps Taxol® precursors, taxadiene, taxa-4(20),11-dien-5α-ol (T5α-ol), and taxa-4(20),11-dien-5-α-yl acetate (T5αAc), via the mevalonate pathway. To screen single genomic modifications, the strains containing single gene deletions or integrations were cultured for three days in galactose or glucose-containing CSM media as simulated previously. In addition to the Taxol® precursors, some side compounds such as verticillene, iso-taxadiene, geranylgeraniol (GGOH), 5(12)-oxa-3(11)-cyclotaxane (OCT), iso-OCT, and additional diterpenes were also detected by GC/MS analyses as previously reported in more detail [20]. Among them, GGOH and OCT were also quantified to compare the production of these side products. Two Taxol® precursors (taxadiene and T5α-ol), two side-products (GGOH and OCT) and biomass were considered in a micro-scale screening in 2 mL cultures to characterize the modified strains. Figure 6 shows the comparison of the production of these metabolites and biomass between the parent strains and designed strains.

**Figure 6:**
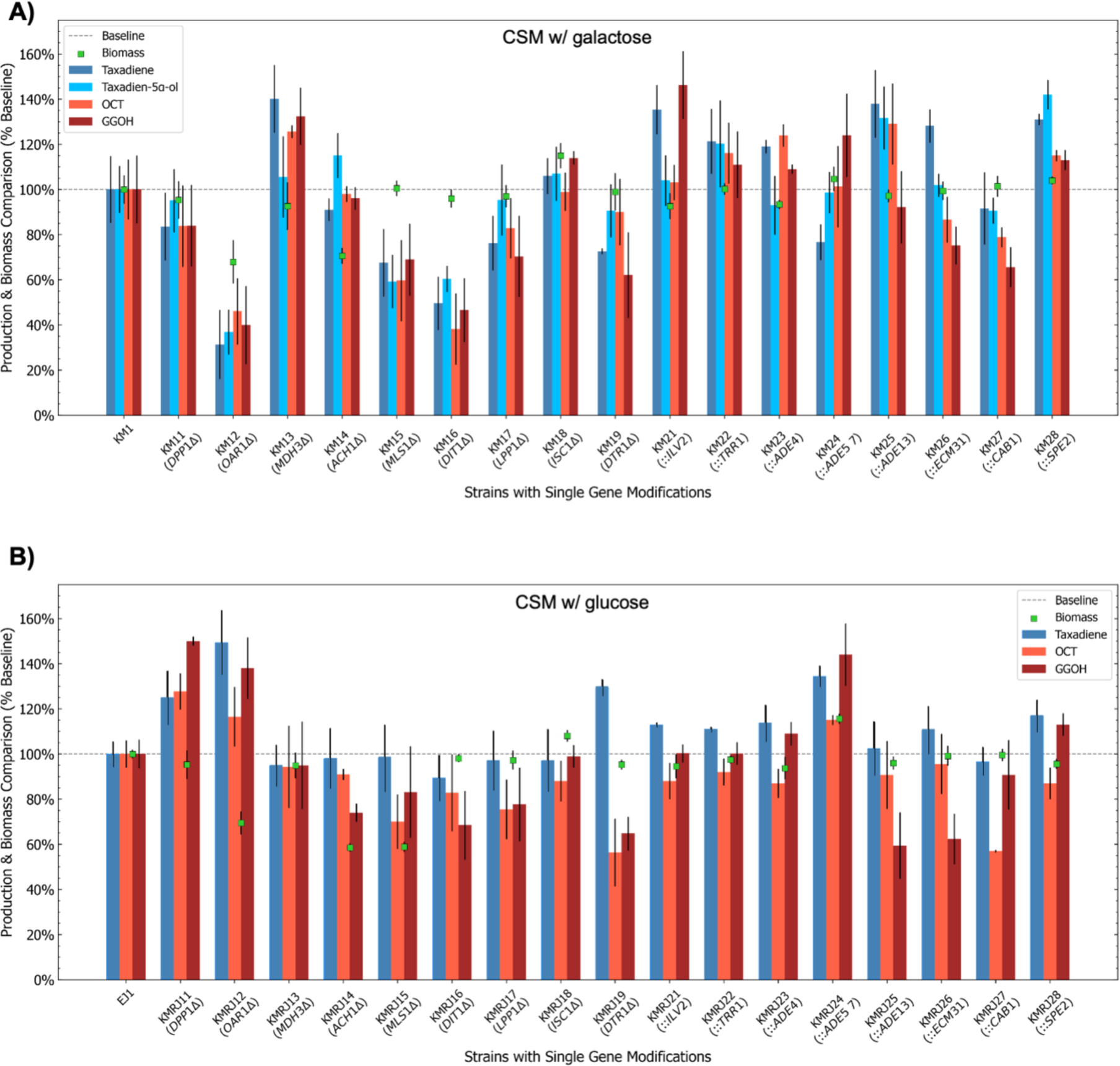
Comparison of biomass and production of early step Taxol® precursors and side products. **A)** KM1-derived strains in the galactose-containing CSM. **B)** EJ1-derived strains in the glucose-containing CSM. Relative values are shown considering 100% baseline (dashed line) of the parent strains, KM1 or EJ1. Error bars indicate standard deviations between three replicates. Δ; gene deletion, ::; gene integration

Differences in production between galactose-containing (CSM w/ galactose) and glucose-containing (CSM w/ glucose) media were detected. Taxadien-5α-ol productions varying from ∼4 mg/L to ∼8 mg/L were observed in galactose-containing media with KM1-derived strains; however, only weak peaks were seen in the gas chromatogram for T5α-ol when the glucose-containing media was used with EJ1-derived strains that did not contain the *GAL80* gene. This was most probably because the expression rates of the galactose-inducible promoters were still higher in galactose-containing media, even though EJ1-derived strains could produce taxadiene under glucose only conditions (Figure 6). Additional regulatory elements might have had an impact on the activation of galactose-inducible promoters in the presence of glucose. In the galactose-containing CSM media, mainly integrations increased the downstream production from GGPP, whereas most of the deletions and integrations showed positive impacts on the production of the precursors in glucose-containing media.

*DPP1* and *LPP1* genes, respectively encode diacylglycerol diphosphate (DGPP) phosphatase and lipid phosphate phosphatase, two Mg^+2^-independent phosphatidate phosphatases found in yeast [46]. As these enzymes are also responsible for converting farnesyl diphosphate (FPP) to farnesol (Figure 3A), they were previously targeted to manipulate the fluxes in the mevalonate pathway to produce plant-sourced sesquiterpenes [47–49]. In the galactose-containing medium, both single deletions showed a similar pattern with lower average (*p*>0.05) taxadiene production (Figure 6A). It could be that the increase in FPP concentration could not make a substantial difference in precursor synthesis as the system was fully active under galactose induction, and previously integrated genes (*ERG20* and *CrtE,* Figure 3A) were able to enhance the flux towards and from FPP sufficiently. On the other hand, a ∼20% increase (*p*<0.05) in taxadiene production with *DPP1* deletion was observed in the glucose-containing medium (Figure 6B), while *LPP1* deletion resulted in almost the same taxadiene production (*p*>0.05). It was reported that *DPP1* deletion could decrease farnesol production [50], and this might be the reason for the higher performance of the *DPP1* deletion (Figure 6B). No significant growth phenotype was observed in any condition (Figure 6). *in silico* simulations suggested that *DPP1* deletion could lead to acetyl-CoA overproduction in a glucose-containing medium (Table 1 and Figure 4B), and experimental results (Figure 6B) supported *in silico* simulations.

The protein encoded by *OAR1* is an NADPH-dependent 3-oxoacyl-(acyl-carrier-protein) reductase, also named 3-keto-acyl ACP reductase, which is involved in fatty acid biosynthesis in yeast [51, 52] as illustrated in Figure 3B. *OAR1* deficient yeast strains could survive on fermentable carbon sources like glucose but not on non-fermentable carbon sources such as glycerol [51, 52]. Approximately 30% reduction in biomass was observed in the glucose-containing and the galactose-containing CSMs (Figure 6). Also, the production rate of the Taxol® precursors obtained from KM12 was less than 40% of the parent strain. In contrast, more than 40% increase was noted in glucose-containing CSM using KMJR12 (Figure 6B). The *OAR1* product catalyzes the reduction of 3-ketoacyl-ACP in mitochondrial fatty acid biosynthesis that uses acetyl-CoA as the source [53, 54]. Therefore, deletion of *OAR1* might indirectly increase acetyl-CoA concentration. KMRJ12 achieved the best taxadiene production among the other strains in glucose-containing CSM. In fact, OptGene predicted that *OAR1* deletion could increase the GGPP production in galactose-containing medium, but we found a better production in glucose-containing CSM. Perhaps, more taxadiene was produced per OD in the galactose-containing medium but low biomass in the galactose-containing CSM resulted in lower total taxane production.

Both OptKnock and OptForce found *MDH3* as a good target to increase acetyl-CoA (Table 1). *MDH3* encodes a peroxisomal malate dehydrogenase [55] responsible for the interconversion of malate and oxaloacetate using NAD^+^ or NADH depending on the reaction in the glyoxylate cycle [56, 57]. OptKnock also suggested deleting the *MLS1* gene that encodes another enzyme, malate synthase, involved in the glyoxylate cycle [56]. Malate synthase catalyzes the formation of malate from glyoxylate and acetyl-CoA in either cytosol or peroxisome, depending on the carbon source [58]. Figure 3C demonstrates the glyoxylate cycle and the roles of *MDH3* and *MLS1* genes. Cytosolic and peroxisomal acetyl-CoA is an essential compound in the glyoxylate cycle. However, the glyoxylate cycle is activated when two-carbon compounds such as ethanol and acetate are used as the sole carbon sources or β-oxidation of fatty acids is enabled [58]. Although we used only six-carbon sources, galactose or glucose, we detected significant effects of *MDH3* or *MLS1* deletions in our study, as shown in Figure 6. We did not observe a considerable growth phenotype for *MDH3* or *MLS1* deletions in galactose-containing CSM, whereas *MLS1* deleted strain KMRJ15 produced less biomass, ∼60% of the reference strain, in glucose-containing CSM. A similar growth trend was previously reported when glucose was used [59]. Yet, the growth behavior of an *MLS1* deleted yeast in a galactose-containing medium has not been reported so far. On the other hand, KM13 (*MDH3*1′) showed a significant increase with ∼40% in taxadiene production when galactose was used as carbon source. Even though it was unexpected as *MDH3* might not be functional in the galactose-containing medium, this increase might be caused by an increased acetyl-CoA concentration as suggested by the algorithms. In contrast, *MLS1* deletion resulted in ∼30% lower taxadiene and possibly lower acetyl-CoA concentrations in KM15. These results might indicate that the glyoxylate cycle genes might still be functional under different carbon sources, and that blocking particular reactions in this cycle can affect cytosolic acetyl-CoA synthesis even if the carbon source is not a two-carbon compound or fatty acid. The FVA analyses indicated that *MDH3* deletion might increase cytosolic acetyl-CoA (Figure 4A-4B); this was in agreement with the experimental results when galactose-containing media was taken into account.

In mitochondria, acetyl-CoA hydrolase (Ach1) encoded by *ACH1* can catalyze the conversion of acetyl-CoA into acetate and CoA. Ach1 can also reversibly transfer CoA from succinyl-CoA to acetate [60]. Blocking these reactions can directly affect acetyl-CoA concentration in the mitochondria, and it might trigger a change in cytosolic acetyl-CoA concentration even though acetyl-CoA cannot be readily transferred from mitochondria to cytosol [60]. OptKnock and OptGene proposed *ACH1* deletion to increase acetyl-CoA concentration in glucose-containing media. Although we detected significantly lower biomass (*p*<0.05) in both KM14 and KMRJ14 strains (Figure 6), taxadiene concentrations were comparable with the reference strains, indicating that production of Taxol® precursors was improved per OD with *ACH1* deletion even if the total production could not be enhanced.

Perhaps, the most surprising findings of OptGene were the deletions of *DIT1* and *DTR1* genes that take part in the sporulation process [61, 62]. Dit1, the product of the *DIT1* gene, plays a role in forming the dityrosine layer of the spore wall by *N*-formylation of tyrosine, and its expression rate increases during spore wall maturation [63]. Dit1 expression might be active in vegetative cells; yet, its cellular activities of are not fully elucidated [64]. Dityrosine transporter encoding gene *DTR1* is also a member of the gene network controlling the assembly of the spore wall [65]. Dtr1 protein can also increase the resistance against some growth inhibitors in vegetative yeast cells [62]. The deletions of these genes did not show a significant decrease in biomass in either glucose-containing or galactose-containing CSM (Figure 6). However, *DIT1* deletion in KM16 caused a dramatically lower production of the Taxol precursors in the galactose-containing medium. In contrast, *DTR1* deletion increased taxadiene concentration in the glucose-containing medium. It is very likely that deletion of these sporulation-specific genes directly makes an impact on GGPP concentration, as it was reported that geranylgeranyl diphosphate synthase has a critical role in the sporulation process in fission yeast, and related genes have functional similarity with *S. cerevisiae* [66]. However, the interaction between *DIT1, DTR1* and the genes involved in GGPP synthesis is not clearly understood yet.

*ISC1* encodes inositol phosphosphingolipase C that hydrolyses sphingolipids to produce ceramide, a compound involved in regulations of cell growth, death and stress response [67]. There is no direct relation between *ISC1* and acetyl-CoA or GGPP, and it was shown that *ISC1* deletion might lead to a higher budding pattern than the wild-type strains [68]. Consistently, *ISC1* deletions resulted in higher average biomass in galactose or glucose-containing media (Figure 6). OptGene suggested *ISC1* deletion to increase acetyl-CoA concentration in a galactose-containing CSM. Parallel to this, we obtained a higher average production for the target metabolites with a ∼6% increase in average in taxadiene and T5α-ol in the galactose-containing medium, although it was not statistically significant (*p*>0.05).

When it comes to the genomic integrations (overexpressions), the majority of the OptForce predictions were involved in the biosynthesis of metabolic precursors in yeast. *ILV2* encodes acetolactate synthase that catalyzes the conversion of two pyruvate molecules to 2-acetolactate in the first steps of isoleucine and valine biosynthesis pathways [69]. From 3-methyl-2-oxobutanoate produced in valine biosynthesis, coenzyme-A precursors are produced in the phosphopantothenate biosynthetic pathway, where *ECM31* and *CAB1* catalyze the first and fourth steps, respectively [70]. The product of *SPE2* plays a critical role in spermidine biosynthesis [71]. At first glance, these genes and pathways might look irrelevant and independent from each other. However, when the intermediate or final products in these pathways are connected as shown in Figure 3D, it becomes clearer that these pathways have direct impacts on CoA production. Therefore, enhanced fluxes in these pathways may also enhance acetyl-CoA production. *ADE4, ADE5,7* and *ADE13* are responsible for five reactions in *de novo* purine synthesis, as shown in Figure 3E [72]. The only exception was the *TRR1* gene, whose product is not a part of a biosynthetic pathway. *TRR1* encodes a regulatory enzyme for the thioredoxin system, a reductase system protecting yeast against reductive or oxidative stress [73].

Indeed, most of the integrations showed an increase in the average production of the target metabolites (Figure 6). Integration of *ADE5,7* encoding a bifunctional protein responsible for the second and fifth reaction in *de novo* purine synthesis pathway resulted in lower taxediene production (∼ 23% decrease, *p*<0.05) in galactose-containing CSM with strain KM24. In contrast, taxediene production was significantly higher in the glucose-containing medium. Apart from this, overexpression of the *CAB1* (coenzyme A biosynthesis 1) gene could not increase the production even though an increased flux in the phosphopantothenate pathway could potentially increase acetyl-CoA concentration in the cell. Recently, Olzhausen et al. (2021) reported that the native *CAB1* is relatively inefficient for the production of coenzyme A compared to other phosphopantothenate pathway genes, and the researchers dramatically increased coenzyme A titer using mutant *CAB1* W331R in rich media [74]. This might be the reason behind non-effective taxane production when *CAB1* was overexpressed. OptForce suggested upregulation of early-stage and mid-stage reactions in *de novo* purine synthesis that have relatively lower fluxes than the late-stage reactions, as shown in Figure S8. This is probably an effective strategy to enhance the flux towards the downstream part of the pathway. Still, the relation of *de novo* purine synthesis pathway and thioredoxin system with acetyl-CoA should be further investigated as increased fluxes in these systems can potentially increase acetyl-CoA concentration or the fluxes in the mevalonate pathway towards GGPP.

These findings show that mathematical modelling and optimization of the microbial systems can be beneficial in finding useful genomic modifications that are very difficult to be intuitively predicted. Indeed, detecting small changes using a relatively low number of replicates is quite challenging as the noise deviations can make the screening and evaluations harder. Nevertheless, statistically significant impacts of the genomic alterations were detected when single modifications were tested. Here, we coupled the native genes with strong galactose-inducible promoters. Therefore, alternative approaches such as protein engineering or integrating heterologous genes encoding higher performance enzymes for the target reaction could also be used to enhance the fluxes in the pathways of interest.

### Next Level Screening using High-throughput Microscale Bioreactor System

To test the effects of multiple modifications on taxane production, the promising modifications were combined to construct *next-level* strains. For galactose-containing CSM, five genomic integrations and one gene deletion were combined to construct different combinations of KM1-derived strains, while three gene deletions and two genomic integrations were used to produce EJ1-derived strains to be used in glucose-containing CSM, as shown in Table 2.

**Table 2:**
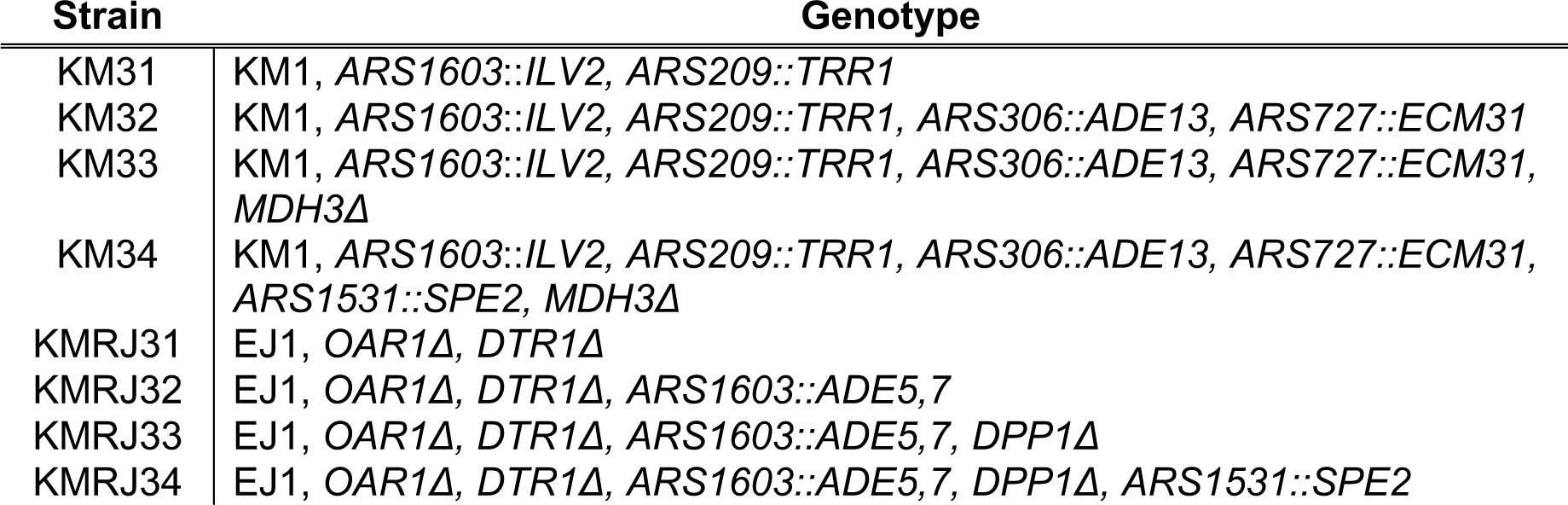
The strains containing multiple modifications

The *first-level* strain screening revealed promising genomic modifications enhancing GGPP and taxadiene productions, as discussed above. However, the next compound, T5α-ol, in the Taxol® pathway could not be effectively detected as a maximum of 8 mg/L production was achieved with KM28 containing an additional copy of the *SPE2* gene (Figure 3D), and only KM25 and KM28 showed a significant increase (*p-* value<0.05) in T5α-ol titer compared to the parent strain (Figure 6). In addition, KMRJ strains could not form a clear T5α-ol peak in the GC chromatogram when they were used in glucose-containing media. The reason behind it was probably the limited agitation and air transfer in deep-well plates because of the shape of the wells and the shaking frequency (350 rpm) as it was proven that oxygen supply is critical for oxygenation reactions for T5α-ol synthesis [20]. Therefore, in the second-level screening, the BioLector Pro microbioreactor-screening platform was utilized with a flower-shaped plate and a higher shaking frequency of 1000 rpm to ensure sufficient oxygen transfer since the flower shape geometry can provide a better oxygen supply [75].

For the galactose-containing medium, the three best-performing strains containing single modifications (KM21, KM25 and KM13) and four strains containing multiple modifications (KM31, KM32, KM33 and KM34) were screened while biomass, pH and DO were monitored in real-time using the BioLector. Interestingly, all multiple modification strains resulted in higher biomass compared to the parent strain, KM1, after three days. It seems increasing fluxes in particular reactions and pathways ended in favor of biomass rather than a metabolic burden on the cell although overexpression of *ILV2* (KM21) or deletion of *MDH3* (KM13) led to lower biomass compared to the parent strain KM1. Most of the strains showed a similar pattern for pH change during three days of cultivation. The initial pH between 5.50 and 5.75 dropped to ∼5.0 and slightly increased to around 5.25, as shown in Figure 7B. *S. cerevisiae* tends to acidify the culture pH [76], therefore it was an expected behavior. However, KM31 containing overexpressed *ILV2* and *TRR1* genes showed a higher pH between 20 and 50 hours of culture. To further elucidate the reason behind this, the pH pattern of KM22 cultures should also be tested.

**Figure 7:**
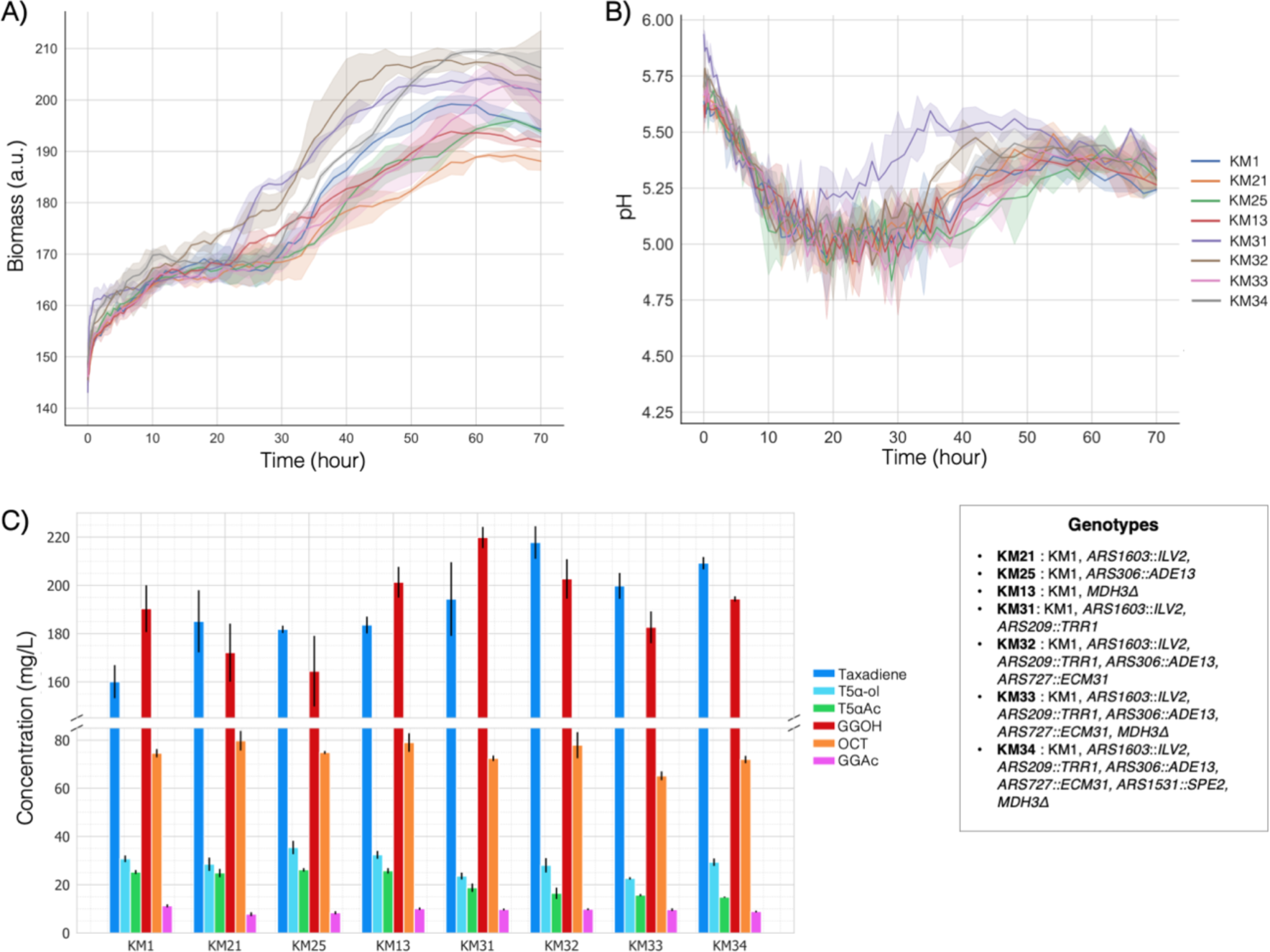
Performances of the selected KM1-derived strains in galactose-containing CSM in the high-throughput microbioreactor system. **A)** Biomass trend **B)** Change in culture pH **C)** Production of the target molecules and side-products Error bars represent the standard deviation of three replicates.

Although KM21, KM25 and KM13 strains increased taxadiene production by 35% - 40% in the deep well-plates compared to KM1 (Figure 6A), taxadiene concentration was only enhanced by ∼15% (*p-*value<0.05) by these strains in the BioLector (Figure 7C). However, the highest taxadiene concentration was ∼110 mg/L in deep-well plates and it was doubled in the BioLector system as ∼215 mg/L production was recorded with KM32. This is a remarkable improvement as approximately 1.6-fold higher production was achieved than the previously reported maximum, which was 137 mg/L in *S. cerevisiae* cell factories [20]. It should also be noted that the previous studies focusing on taxadiene synthesis used richer media that are likely to increase the production yield. In this study, complete synthetic defined media were used since all *in silico* illustrations were conducted using defined media. Even though a higher taxadiene production was achieved with 1 g/L in *E. coli* [77], the expression of the second step enzymes, T5αOH and CPR, totally zeroed the taxadiene synthesis in *E. coli* [77]. Therefore, KM32 increased taxadiene production while it also expressed the downstream genes, T5αOH, CPR and TAT, that are responsible for the two subsequent steps.

Dissolved oxygen (DO) was maintained between 75% - 100% for all the strains used in the BioLector system (Figure S10 and S11). Accordingly, greater T5α-ol was synthesized by all KM1-derived strains with at least 23 mg/L concentration in KM33 (Figure 7C). This production was almost three-fold higher than the maximum T5α-ol production observed in deep-well plates, proving that oxygen supply was the critical factor for T5α-ol production. Also, the acetylated precursor, T5αAc, was detected in all KM1-derived strains with concentrations ranging from 15 to 26 mg/L. Nevertheless, all multi-modification strains KM31, KM32, KM33 and KM34, produced less T5αAc (*p-* value<0.05) than the parent strain as shown in Figure 7C even though a significant increase (*p-*value<0.05) was observed in taxadiene production in KM32, KM33 and KM34 and single-modification strains (KM21, KM25, KM13). This indicates that to enhance the yield of T5α-ol and the following compound, T5αAc, further improvement in the second reaction catalyzed by T5αOH and CPR is a necessity. KM25 was the best producer of T5α-ol and T5αAc with 35.4 mg/L and 26.2 mg/L, respectively. Yet, no statistically significant difference (*p-*value>0.05) was observed for T5αAc titer between KM1, KM21, KM25 and KM13. In addition to taxadiene, these are the highest yields for T5α-ol and T5αAc production reported so far using *S. cerevisiae* as a cell factory. It should be also noted that 23 mg/L T5α-ol was produced by *E. coli* in the previous attempt [77] without the expression of the next step gene, TAT, in the pathway. Also, this *E. coli* strain could not accumulate taxadiene as mentioned above. Considering these, further improvements are likely to increase the flux towards the T5αAc in our *S. cerevisiae* strains.

In the glucose-containing CSM, KMRJ12, KMRJ19 and KMRJ24 strains were used with multi-modification strains KMRJ31, KMRJ32, KMRJ33, and KMRJ34 (Table 2) in the BioLector platform. In parallel to the deep-well plates, all strains with the *OAR1Δ* genotype produced less biomass in the BioLector system as shown in Figure 8A. A correlation was also observed between pH change and biomass and/or *OAR1Δ* (Figure 8B). Like in the galactose-containing medium, after a sharp drop, the pH reached around 5.25 for the strains expressing the *OAR1* gene (EJ1, KMRJ19, KMRJ24). The pH stabilised at around 5.0 for the other strains, this might be because of lower biomass or the *OAR1* deletion.

**Figure 8:**
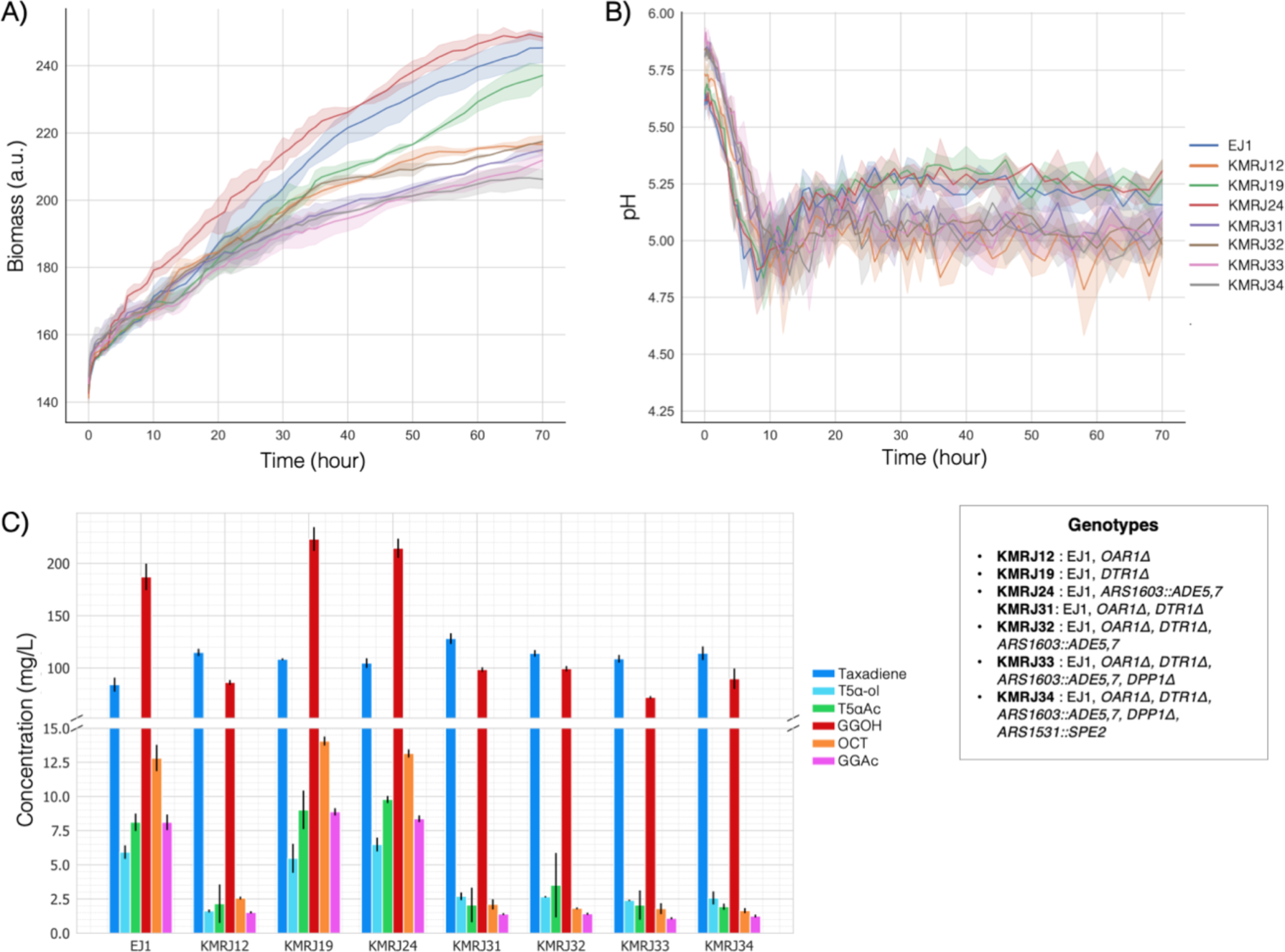
Performances of the selected EJ1-derived strains in glucose-containing CSM in the high-throughput microbioreactor system. **A)** Biomass trend **B)** Change in culture pH **C)** Production of the target molecules and side-products Error bars represent the standard deviation of three replicates.

Another interesting finding with *OAR1* deletion was the dramatic reductions in the titers of the oxygenated product, T5α-ol, and naturally, the next product T5αAc as seen in Figure 8C. Also, GGOH production dropped at least 2-fold in all *OAR1-*deleted strains. GGOH is indeed a valuable compound that can be used in the pharmaceutical and cosmetic industries. Metabolic engineering studies have been carried out to improve GGOH production in *S. cerevisiae* [78, 79]. Although the GGOH synthesis is not fully identified in yeast, native phosphatases are thought to be involved in GGOH synthesis from GGPP. Considering its chemical structure, oxygenation reactions should also take a part in GGOH synthesis [80]. Taking the reductions in T5α-ol and GGOH into consideration, it is likely that *OAR1* deletion affected the oxygenation capability of *S. cerevisiae,* and it might also be one of the reasons behind the decrease in cell fitness. On the other hand, still, the strains with the *OAR1Δ* genotype showed greater taxadiene production than the reference strain (Figure 8C), meaning that *OAR1* deletion led to the highest taxadiene production per OD. However, it should be also noted that GGOH reduction might have improved the taxadiene concentration.

The combination of *OAR1Δ* and *DTR1Δ* in KMRJ31 resulted in the maximum taxadiene production of 128 mg/L, which is still a remarkable titer for taxadiene considering the similar studies reported previously [19, 81]. On the other hand, even if *OAR1Δ* genotypes were excluded, KMRJ strains produced ∼7 mg/L of T5α-ol from each 100 mg/L taxadiene on average (7% yield), whereas this amount was ∼18 mg/L for KM strains (18% yield). This is probably because the galactose-inducible promoters could not express T5αOH in glucose-containing CSM as efficiently as in galactose-containing CSM, even though the *GAL80* was deleted.

In addition to Taxol® precursors, KMRJ19 with *DTR1* deletion produced the highest GGOH with 221 mg/L among all the strains used in this study, it was followed by KMRJ24 and KM31in the galactose-containing medium with the 219 mg/L (*p-* value>0.05). These production yields of GGOH are comparable with those in similar studies [78, 79]. For this reason, the related modifications, *DTR1* deletion*, ADE 5,7* overexpression or simultaneous overexpression of *ILV2* and *TRR1*, are promising genomic modifications to increase GGOH production.

### Scale-up of the Production of the Taxol® Precursors using the best strain

Finally, the production was scaled-up using the best taxadiene producer, KM32, in 250 mL reaction volume in a mini-scale bioreactor. To maintain similar conditions as in the BioLector system, minimum %DO was set to 75% and O_2_ was supplied with air when it was below the threshold. Likewise, pH was set to 6.0 as it is a suitable pH considering both yeast growth and the enzymatic activities in the Taxol® pathway [20]. A 1M NaOH was added as needed when the pH was less than 6.0. Figure 9 shows the reactor parameters and the production of Taxol® precursors during the five days of cultivation.

**Figure 9.**
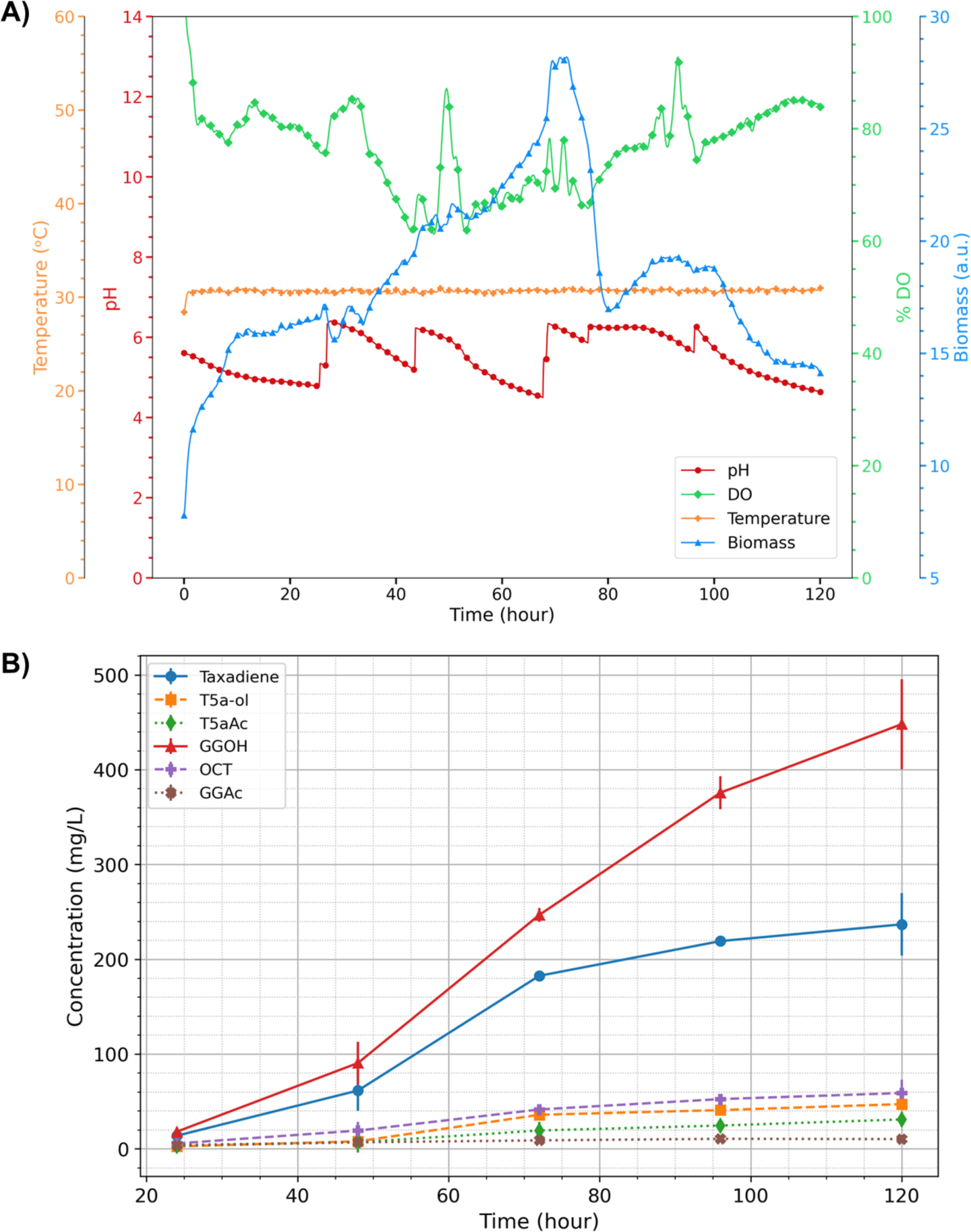
**A)** The trends of the bioreactor parameters in KM32 culture **B)** Metabolite concentrations during five days of cultivation. Error bars represent the standard deviation of two measurements taken each time. Total working volume was set to 250 mL: 200 mL of the culture and 50 mL of the dodecane layer.

Considering the first three days, the growth showed a similar pattern as a decrease was observed towards the third day and then dramatically dropped down (Figure 9A). Although this decrease was expected, the sharp drop between 72^nd^ and 80^th^ hours was probably a temporary a sensor problem. On the other hand, the production yield was relatively low for the Taxol® precursors in the first three days. In 72 hours, 182.5 mg/L taxadiene, 35.8 mg/L of T5α-ol and 19.2 mg/L of T5αAc were produced. These reached to 236.7, 47.1 and 31.2 mg/L, respectively on the fifth day (Figure 9B). Although the total titers of the target molecules were higher at the end of cultivation, on the fifth day, the yield on the third day was lower compared to the BioLector system. On the other hand, GGOH concentration was dramatically higher than the taxadiene concentration (Figure 9B). pH and DO showed more fluctuations in the bioreactor-scale. This might be the reason behind the decrease in production yield; therefore, an optimized bioprocess condition can further improve the titers of Taxol® precursors in larger scales.

### Comparison of the *in silico* Predictions and Wet-lab Results

The wet-lab validations showed that most of the genomic modifications resulted in higher taxadiene production (and most probably higher GGPP production), highlighting that mathematical models and computational frameworks can help us to find useful targets that might be very difficult to predict intuitively. Considering Table 1, Figure 4, and the experimental results, however, there are also exceptions. *OAR1* deletion was predicted to increase GGPP concentration in the galactose-containing CSM, and *MDH3* deletion was suggested to increase acetyl-CoA production in the cytosol in the glucose-containing medium. Nevertheless, these deletions showed higher productions in the opposite carbon source (Figure 6). According to the *in silico* predictions, *MLS1* deletion should have increased the acetyl-CoA production in the galactose-containing medium, but in contrast, this deletion decreased GGPP concentration and probably acetyl-CoA concentration in the galactose-containing medium and could not make any difference in glucose-containing CSM (Figure 6). The other deletions showed either predicted results with overproductions or similar production yields to the reference strains (*p-*value>0.05). It is also possible that more replicates could have given more precise results and could have mitigated the analytic measurement deviations.

According to these, *in silico* predictions presented a narrowed solution set compared to thousands of gene targets. Still, these predicted genomic modifications should be validated with wet-lab studies. In addition, our findings also showed that the medium conditions, primarily the carbon source, directly affect the performances of the designed strains; therefore, a genomic modification might show a better result in different conditions.

## CONCLUSION

Computational tools, *in silico* design frameworks and genome-scale models were used to rationally design yeast cell factories in a way that intuitive estimations are very difficult. In specific, in this study, a combinatorial design approach was used with three *in silico* algorithms, OptKnock, OptGene and OptForce, on the yeast-GEM 8.5.0. A set of 17 genomic modifications were predicted by the simulations (nine gene knock-outs, eight gene overexpressions and their combinations via extra copies and strong promoters) and tested for the increased production of Taxol® precursors, through GGPP overproduction. Therefore, this study is one of the most comprehensive studies reported until now in terms of the amount and the variability of the genomic manipulations. Without a doubt, the ACtivE toolkit and method facilitated this process with an accelerated genome editing process. The findings of this work showed that simultaneous overexpression of four genes, *ILV2, TRR1, ADE13* and *ECM31,* related to upregulations in the pantothenate pathway, branched-amino acid biosynthesis pathways and *de novo* purine synthesis pathway, could enhance taxadiene production by ∼50% through GGPP overproduction. Also, single modifications could increase taxadiene yield from 15% to 40% depending on the cultivation conditions and carbon source. As the oxygen supply is crucial for the second step in the Taxol® biosynthesis pathway, higher yields in T5α-ol and T5αAc productions were reported with the conditions allowing better oxygen access. Using the best-performing strain KM32 containing additional copies of the above-mentioned four genes, we achieved 215 mg/L of taxadiene, 43.65 mg/L of T5α-ol and 26.2 mg/L of T5αAc titers that are the highest production yields reported until now in *S. cerevisiae.* The genomic modifications reporting higher GGPP synthesis can be also used to increase the production of other high-value isoprenoids through the same precursor. In addition to the mevalonate pathway, similar integrated approaches combining different design algorithms, genome engineering and bioprocessing studies can be used for enhancing metabolic fluxes towards the target native or heterologous pathways in *S. cerevisiae.* Developing alternative *in silico* prediction and design tools, constructing more accurate genome-scale models would improve the efficiency of this process.

## Author contributions

Koray Malcı: Conceptualization, Methodology, Software, Formal Analysis, Investigation, Writing – Original Draft. Rodrigo Santibáñez: Methodology, Software, Writing – Reviewing and Editing. Nestor Jonguitud-Borrego: Investigation, Resources. Jorge Santoyo-Garcia: Investigation. Eduard J. Kherkoven: Supervision, Writing – Reviewing and Editing. Leonardo Rios-Solis: Conceptualization, Supervision, Writing – Reviewing and Editing, Project administration.

## Data/Code availability

MATLAB scripts are available on https://github.com/kmalci/yeast_modelling. Plasmids and other data will be made available on request.

## Acknowledgements

We would like to thank Mr Stuart Martin, Mr Mark Lauchlan and Miss Katalin Kis at the School of Engineering, University of Edinburgh, UK for their technical support with GC-MS analysis and laboratory operations. We would like to thank Dr David Gomez-Cabeza at the School of Engineering, University of Edinburgh, UK for their kind assistance with the inverted microscope.

This work was supported by the YLSY program of the Ministry of National Education of Turkey, the Mexican government’s dependence CONACyT (Mexican National Council for Science and Technology) CVU:675492 and CVU: 537962, Novo Nordisk Foundation (Grant Number NNF20CC0035580), the Engineering and Physical Sciences Research Council (Grant number EP/R513209/1), the Royal Society (Grant Number RSG\R1\180345) and the British Council (Grant Number: 527429894).

## Supplementary data

Lists of primers and DNA parts; genotypes of the strains developed; predicted genomic modifications that were eliminated and not tested experimentally; acetyl-CoA-centered sub-metabolite interaction maps for the selected gene deletions; Escher maps of the metabolic pathways targeted in the study; DO trends measured in the Biolector experiments; gas chromatography results showing the production of Taxol® precursors.

