## Supplemental Data for "Improved Production of Taxol® Precursors in *S. cerevisiae* using Combinatorial *in silico* Design and Metabolic Engineering"

**Table S1:** The primer list used to produce the donor DNA parts for the gene deletions

| **Name** | **Sequence 5' to 3'** |
|---|---|
| DPP1 UHA For | CAAGTTCGTTGCACTGTATTTC |
| DPP1 UHA Rev + DHA | AGAATCAGAATTAAATCATAGCAAACGACCCCACATGACATACGAAATATACG |
| DPP1 DHA For + UHA | ATAAATACGTATATTTCGTATGTCATGTGGGGTCGTTTGCTATGATTTAATTC |
| DPP1 DHA Rev | GTATCAGTCACAGGTACGG |
| DPP1 Col Rev | CCATTGATGATCCTCTTCCG |
| OAR1 UHA For | CTTCAACATCATCGTCACC |
| OAR1 UHA Rev + DHA | TGGTATAGACGTGGGAGAAGAAAAGTCTGGCAGTGAACACATTTTCACATC |
| OAR1 DHA For + UHA | AAGAAACGTGATGTGAAAATGTGTTCACTGCCAGACTTTTCTTCTCCCAC |
| OAR1 Col Rev | GATTTACAGAAGTTTTAGCTGC |
| MDH3 UHA For | CAACACATCGGTCATAGTAG |
| MDH3 UHA Rev + DHA | CTTCGTAAATCAAGGGAAAACACTTGTCAGGAAGGAAAGGAAACCATATCC |
| MDH3 DHA For + UHA | CCACTAGGAGGATATGGTTTCCTTTCCTTCCTGACAAGTGTTTTCCCTTG |
| MDH3 DHA Rev | CTCTCAGTTGATTTTCCGTG |
| MDH3 Col Rev | GTATCCATCGATACCAGTGT |
| ACH1 UHA For | CATATGACATACGTATTAGCCGC |
| ACH1 UHA Rev + DHA | ATAAATATGCAAGAAAAAACAACGCATTGGGTTTGTTTTGCCGCTATTGTC |
| ACH1 DHA For + UHA | GTCTTACTAGACAATAGCGGCAAAACAAACCCAATGCGTTGTTTTTTCTTG |
| ACH1 DHA Rev | CACCACTGATAGATAACACACA |
| ACH1 Col Rev | GAGTACCCACTCCTGTAAAAC |
| MLS1 UHA For | CAGTATTACCCTACATTGCTATC |
| MLS1 UHA R + DHA | AGATGATTCATTGCTAACTACGAAACGAAGGTGCTTTTACTACTTTGTTTAGTTC |
| MLS1 DHA For + UHA | GTTTTGAACTAAACAAAGTAGTAAAAGCACCTTCGTTTCGTAGTTAGCAATG |
| MLS1 DHA Rev | CACACTCATATGTGAGTTAGATG |
| MLS1 Col Rev | GGTAAGAATATCTCCCACTGTAG |
| DIT1 UHA For | GAGTCCCTGGAAGGAAAATTATTG |
| DIT1 UHA Rev + DHA | TATTCCCCCTCTGTTAAATGGAATTGTGTGGCGGAGGAGCACAATTTATG |
| DIT1 DHA For + UHA | TAAATTTTCACATAAATTGTGCTCCTCCGCCACACAATTCCATTTAACAGAGG |
| DIT1 DHA Rev | CTGATGCCTCAAGATTTAACC |
| DIT Col Rev | GATGAAGAAAATGAAGAGGCAGG |
| LPP1 UHA For | CACCGACGGATTCAGAG |

| | |
|---|---|
| LPP1 UHA Rev + DHA | CATCAACGCCTAAGGAAACTCGTCATATTCCACTTACAGAGTCCTATCAGG |
| LPP1 DHA For + UHA | TTATTCTTTCCTGATAGGACTCTGTAAGTGGAATATGACGAGTTTCCTTAGG |
| LPP1 DHA Rev | CTATATGAAAACTTCACAGGGAG |
| LPP1 Col Rev | GTCTCTGGAGCTGTTCTAG |
| ISC1 UHA For | CGTAATTGGACACACTCTTTAC |
| ISC1 UHA Rev + DHA | TTCTCCGTGATTGCTTTGCATCTATTGACGCTCGAAGATTTAAGCCAAACC |
| ISC1 DHA For + UHA | ATTTATTTTGGTTTGGCTTAAATCTTCGAGCGTCAATAGATGCAAAGCAATC |
| ISC1 DHA Rev | GACATTTACTTAATGTGGAGCAC |
| ISC1 Col Rev | CTTCAAACTCAGATTGACCATC |
| DTR1 UHA For | GTTGCTATGTTCGGATGTAC |
| DTR1 UHA Rev + DHA | TTTCTTGAAGTCCTTGGGATGAGTGATGAGCAACAACAACTTTTCTTACTACC |
| DTR1 DHA For + UHA | GAAGGATGGTAGTAAGAAAAGTTGTTGTTGCTCATCACTCATCCCAAGGAC |
| DTR1 DHA Rev | GGTCTAACTGACACTACTGTTCC |

* The **red** sequences show the overlapping fragments with neighbour parts, while the **black** sequences show the annealing fragments.
** Plus (+) sign shows the overlapping neighbour parts targeted by the **red** sequences
*** If the overlapping sequences (**red**) show higher affinity than the annealing sequences (**black**) for the same DNA template, only annealing parts should be used first to amplify the target regions.
**** For: forward, Rev: reverse, UHA: upstream homology arm, DHA: downstream homology arm, Col: for colony PCR (coupled with corresponding UHA For)

**Table S2:** crRNA sequences used to target the corresponding genes for deletions

| Target Gene | Sequence 5' – 3' | Chromosome |
|---|---|---|
| *DPP1* | GCAGTAAATAAAGTGTCCAA *TGG* | IV |
| *OAR1* | AGGAGATGGCCGGAACTCAG *TGG* | XI |
| *MDH3* | TGTTATTGGGGGTCATTCAG *GGG* | IV |
| *ACH1* | CTGAAAGTGGACGACAAGTG *TGG* | II |
| *MLS1* | AAGATTACATTGGGATCCCA *AGG* | XIV |
| *DIT1* | CCTTGTCAAGATATATCCAC *CGG* | IV |
| *LPP1* | ATGAAACTTGAATGTCCGCT *TGG* | IV |
| *ISC1* | ACTCCTGGGAGCAATTGCAT *GGG* | V |
| *DTR1* | GGTAATGGAGACCCTAAATG *GGG* | II |

* The **red** nucleotides represent the corresponding PAM sequences

**Table S3:** The primer list used to produce the donor DNA parts for the genomic integrations

| Name | Sequence 5' to 3' |
|---|---|
| UHA 1603 For | GGCTATGGTGGTGATGTCTG |
| DHA 1603 Rev | CTGTCTCCGCTATGTCAGTTAC |
| UHA 1603 Rev + pGAL1 | CTAATCCGTACTTCAATATAGCAATGAGCGAGGAAAAAAACAGTTGTAC ATTGG |
| pGAL For + UHA 1603 | GTTACCAATGTACAACTGTTTTTTTCCTCGCTCATTGCTATATTGAAGTA C |
| pGAL Rev + ILV2 | GAAGTTTTTTAGCGTAGATTGTCTGATCATTATAGTTTTTTCTCCTTGACG |
| ILV2 For + pGAL | ATACTTTAACGTCAAGGAGAAAAAACTATAATGATCAGACAATCTACGCTA AAAAAC |
| ILV2 Rev + DHA 1603 | AAAAGCTCGTGAATACGGCAAGAACGAAGCGCGGTTAAATTCGTATTGG CC |
| DHA 1603 For + ILV2 | AAGAACAGTGGCCAATACGAATTTAACCGCGCTTCGTTCTTGCCGTATTC |
| pGAL Rev + TRR1 | ACCAATGATAGTAACTTTGTTGTGAACCATTATAGTTTTTTCTCCTTGACG |
| TRR1 For + pGAL | ATACTTTAACGTCAAGGAGAAAAAACTATAATGGTTCACAACAAAGTTACT ATC |
| TRR1 Rev + DHA 1603 | AAAAGCTCGTGAATACGGCAAGAACGAAGCGTCAGTAGCTGTAATATCA GATG |
| DHA 1603 For + TRR1 | AATAAAGCATCTGATATTACAGCTACTGACGCTTCGTTCTTGCCGTATTC |
| pGAL Rev + ADE4 | TGCTAATACAATACCTAAAATACCACACATTATAGTTTTTTCTCCTTGACG |
| ADE4 For + pGAL | ATACTTTAACGTCAAGGAGAAAAAACTATAATGTGTGGTATTTTAGGTATT G |
| ADE4 Rev + DHA 1603 | AAAAGCTCGTGAATACGGCAAGAACGAAGCCTTTTCTTTTTGTACAAGCT G |
| DHA 1603 For + ADE4 | AGCAATCCGCAGCTTGTACAAAAAGAAAAGGCTTCGTTCTTGCCGTATTC |
| pGAL Rev + ADE57 | ACCGTTTCCTAAAACGAGAATGTTGAGCATTATAGTTTTTTCTCCTTGACG |
| ADE57 For + pGAL | ATACTTTAACGTCAAGGAGAAAAAACTATAATGCTCAACATTCTCGTTTTA GG |
| ADE57 Rev + DHA 1603 | AAAAGCTCGTGAATACGGCAAGAACGAAGCCTACAACTCCTCAACAGTAT TCTC |
| DHA 1603 For + ADE57 | TAGCAAGAGAATACTGTTGAGGAGTTGTAGGCTTCGTTCTTGCCGTATTC |
| pGAL Rev + ADE13 | TGGCGTAGTGTAATTGTCGTAGTCAGGCATTATAGTTTTTTCTCCTTGACG |
| ADE13 For + pGAL | ATACTTTAACGTCAAGGAGAAAAAACTATAATGCCTGACTACGACAATTAC |
| ADE13 Rev + DHA 1603 | AAAAGCTCGTGAATACGGCAAGAACGAAGCCGACTGAGTAAGAACCGTT TC |
| DHA 1603 For + ADE13 | TAAAAATATGAAACGGTTCTTACTCAGTCGGCTTCGTTCTTGCCGTATTC |
| pGAL Rev + ECM31 | GGTGCATAATTGTCTTTTCATTATATTCATTATAGTTTTTTCTCCTTGACG |
| ECM31 For + pGAL | ATACTTTAACGTCAAGGAGAAAAAACTATAATGAATATAATGAAAAGACAA TTATGC |
| ECM31 Rev + DHA 1603 | AAAAGCTCGTGAATACGGCAAGAACGAAGCGATAAAGACAACAACTCGT G |
| DHA 1603 For + ECM31 | CTTGGTAGCCCACGAGTTGTTGTCTTTATCGCTTCGTTCTTGCCGTATTC |
| pGAL Rev + CAB1 | GTAAGATATCTCTTGAGTAATTCGCGGCATTATAGTTTTTTCTCCTTGACG |

| | |
|---|---|
| CAB1 For + pGAL | ATACTTTAACGTCAAGGAGAAAAAACTATAATGCCGCGAATTACTCAAGAG |
| CAB1 Rev + DHA 1603 | AAAAGCTCGTGAATACGGCAAGAACGAAGCCCCAGCTTGGAAAAGTACAATTTG |
| DHA 1603 For + CAB1 | TTATTTCAAATTGTACTTTTCCAAGCTGGGGCTTCGTTCTTGCCGTATTC |
| pGAL Rev + SPE2 | GTTAGTCAATTCTTTTATGGTGACAGTCATTATAGTTTTTTCTCCTTGACG |
| SPE2 For + pGAL | ATACTTTAACGTCAAGGAGAAAAAACTATAATGACTGTCACCATAAAAGAATTG |
| SPE2 Rev + DHA 1603 | AAAAGCTCGTGAATACGGCAAGAACGAAGCGATATTAAATTAGCGTGCTTGC |
| DHA 1603 For + SPE2 | CCCTCCAAGCAAGCACGCTAATTTAATATCGCTTCGTTCTTGCCGTATTC |
| ILV2 Col Rev | GCGGTATGCTATATGTTGAAAGC |
| TRR1 Col Rev | CTTTATTGTTCGGAGCAGTGC |
| ADE4 Col Rev | CATCTTGTCCACGATGTTGTAG |
| ADE57 Col Rev | CTTGGTGACAAGAACGTGTTC |
| ADE13 Col Rev | GTTGCTGACATTTCCTTGGAG |
| ECM31 Col Rev | CATACGCAGTACACATCGACA |
| CAB1 Col Rev | GCAAGGTTGAAAGTATTGTCGC |
| SPE2 Col Rev | GCTGATAGTTCGTGGTCAATG |
| 209 UHA For | GAATGTCCGTGGTAATACAATGG |
| 209 UHA Rev + pGAL | CTAATCCGTACTTCAATATAGCAATGAGCCTAGCACATTTTATGGGCCTAAG |
| pGAL For + 209 UHA | ATAATGTCTTAGGCCCATAAAATGTGCTAGGCTCATTGCTATATTGAAGTAC |
| TRR1 Rev + 209 DHA | TTGAATACAGAGCAAAAGGATTAGCCATACGTCAGTAGCTGTAATATCAGATG |
| 209 DHA For + TRR1 | AATAAAGCATCTGATATTACAGCTACTGACGTATGGCTAATCCTTTTGCTCTG |
| 209 DHA Rev | CTCTATATCGCTGTTGCTTATGG |
| 306 UHA For | GTGACTGTCTCCAAGAATACGAC |
| 306 UHA Rev + pGAL | CTAATCCGTACTTCAATATAGCAATGAGCGCTCCTTCTCCTAACATCAATAACG |
| pGAL For + 306 UHA | CTGTTCGTTATTGATGTTAGGAGAAGGAGCGCTCATTGCTATATTGAAGTAC |
| ADE13 Rev + 306 DHA | TTCAGAAACACTGCTTACACTATTCACCAGCGACTGAGTAAGAACCGTTTC |
| 306 DHA For + ADE13 | TAAAAATATGAAACGGTTCTTACTCAGTCGCTGGTGAATAGTGTAAGCAGTGTTTC |
| 306 DHA Rev | CAAGAACACCAGACCTCCAAGC |
| 727 UHA For | GACTTGGAAGACCACACTACTCTCC |
| 727 UHA Rev + pGAL | CTAATCCGTACTTCAATATAGCAATGAGCCATAGCAGTGGCGCGGTC |
| pGAL For + 727 UHA | TTATAGGGAATCGACCGCGCCACTGCTATGGCTCATTGCTATATTGAAGTAC |
| ECM31 Rev + 727 DHA | ATCAGCAGGCCATGGATAAACTTTCCGTTGGATAAAGACAACAACTCGTG |
| 727 DHA For + ECM31 | CTTGGTAGCCCACGAGTTGTTGTCTTTATCCAACGGAAAGTTTATCCATGG |
| 727 DHA Rev | GAGATTCTTGACGTAAAGTGC |

| | |
|---|---|
| 1306 UHA For | GGTTTCAAGCCAAATTGTACG |
| 1306 UHA Rev + pGAL | CTAATCCGTACTTCAATATAGCAATGAGCCTTAGGTAGTAACTATACGCAGC |
| pGAL For + 1306 UHA | GGAGCAGCTGCGTATAGTTACTACCTAAGGCTCATTGCTATATTGAAGTAC |
| ADE57 Rev + 1306 DHA | TAGCCCACTTCTAGCCAACTTCTAGCCCACCTACAACTCCTCAACAGTATTCTC |
| 1306 DHA For + ADE57 | TAGCAAGAGAATACTGTTGAGGAGTTGTAGGTGGGCTAGAAGTTGGCTAGAAG |
| 1306 DHA Rev | GCGCATAGTGCTAGTCTTTTCTCC |

* The **red** sequences show the overlapping fragments with neighbour parts, while the **black** sequences show the annealing fragments.
** Plus (+) sign shows the overlapping neighbour parts targeted by the **red** sequences
*** If the overlapping sequences (**red**) show higher affinity than the annealing sequences (**black**) for the same DNA template, only annealing parts should be used first to amplify the target regions.
**** For: forward, Rev: reverse, UHA: upstream homology arm, DHA: downstream homology arm, Col: for colony PCR (coupled with corresponding UHA For)

**Table S4:** The yeast strains used in the study

| Strain | Genotype | Source |
|---|---|---|
| LRS6 | *MATa, leu2-3, 112::HIS3MX6-GAL1p-ERG19/GAL10p-ERG8; ura3-52::URA3-GAL1p-MvaS$^{A110G}$/GAL10p-MvaE; his3Δ1::hphMX4-GAL1p-ERG12/GAL10p-IDI1; trp1-289::TRP1_GAL1p-CrtE (X.dendrorhous)/GAL10p-ERG20; YPRCdelta15::NatMX-GAL1p-CrtE/GAL10p-CrtE*; *ARS1014::GAL1p-TASY-GFP*; *ARS1622b::GAL1p-MBP-TASY-ERG20*; *ARS1114a::TDH3p-MBP-TASY-ERG20; ARS511b::GAL1p-T5αOH/GAL3-CPR; RKC3::GAL1p-TAT* | Walls et al., 2020 |
| KM1 | LRS6, *RKC4*::*GAL1p-TAT* | This study |
| KM11 | KM1, *DPP1Δ* | This study |
| KM12 | KM1, *OAR1Δ* | This study |
| KM13 | KM1, *MDH3Δ* | This study |
| KM14 | KM1, *ACH1Δ* | This study |
| KM15 | KM1, *MLS1Δ* | This study |
| KM16 | KM1, *DIT1Δ* | This study |
| KM17 | KM1, *LPP1Δ* | This study |
| KM18 | KM1, *ISC1Δ* | This study |
| KM19 | KM1, *DTR1Δ* | This study |
| KM21 | KM1, *ARS1603::GAL1p-ILV2* | This study |
| KM22 | KM1, *ARS1603::GAL1p-TRR1* | This study |

| | | |
|---|---|---|
| KM23 | KM1, *ARS1603::GAL1p-ADE4* | This study |
| KM24 | KM1, *ARS1603::GAL1p-ADE5,7* | This study |
| KM25 | KM1, *ARS1603::GAL1p-ADE13* | This study |
| KM26 | KM1, *ARS1603::GAL1p-ECM31* | This study |
| KM27 | KM1, *ARS1603::GAL1p-CAB1* | This study |
| KM28 | KM1, *ARS1603::GAL1p-SPE2* | This study |
| KM31 | KM1, *ARS1603*::*ILV2, ARS209::TRR1* | This study |
| KM32 | KM1, *ARS1603*::*ILV2, ARS209::TRR1, ARS306::ADE13, ARS727::ECM31* | This study |
| KM33 | KM1, *ARS1603*::*ILV2, ARS209::TRR1, ARS306::ADE13, ARS727::ECM31, MDH3Δ* | This study |
| KM34 | KM1, *ARS1603*::*ILV2, ARS209::TRR1, ARS306::ADE13, ARS727::ECM31, ARS1531::SPE2, MDH3Δ* | This study |
| EJ1 | KM1, *GAL80Δ* | This study |
| KMRJ11 | EJ1, *DPP1Δ* | This study |
| KMRJ12 | EJ1, *OAR1Δ* | This study |
| KMRJ13 | EJ1, *MDH3Δ* | This study |
| KMRJ14 | EJ1, *ACH1Δ* | This study |
| KMRJ15 | EJ1, *MLS1Δ* | This study |
| KMRJ16 | EJ1, *DIT1Δ* | This study |
| KMRJ17 | EJ1, *LPP1Δ* | This study |
| KMRJ18 | EJ1, *ISC1Δ* | This study |
| KMRJ19 | EJ1, *DTR1Δ* | This study |
| KMRJ21 | EJ1, *ARS1603::GAL1p-ILV2* | This study |
| KMRJ22 | EJ1, *ARS1603::GAL1p-TRR1* | This study |
| KMRJ23 | EJ1, *ARS1603::GAL1p-ADE4* | This study |
| KMRJ24 | EJ1, *ARS1603::GAL1p-ADE5,7* | This study |
| KMRJ25 | EJ1, *ARS1603::GAL1p-ADE13* | This study |
| KMRJ26 | EJ1, *ARS1603::GAL1p-ECM31* | This study |
| KMRJ27 | EJ1, *ARS1603::GAL1p-CAB1* | This study |
| KMRJ28 | EJ1, *ARS1603::GAL1p-SPE2* | This study |
| KMRJ31 | EJ1, *OAR1Δ, DTR1Δ* | This study |
| KMRJ32 | EJ1, *OAR1Δ, DTR1Δ, ARS1603::ADE5,7* | This study |
| KMRJ33 | EJ1, *OAR1Δ, DTR1Δ, ARS1603::ADE5,7, DPPΔ* | This study |
| KMRJ34 | EJ1, *OAR1Δ, DTR1Δ, ARS1603::ADE5,7, DPPΔ, ARS1531::SPE2* | This study |

**Table S5:** The gene candidates eliminated by *in silico* analyses, FVA and metabolite interaction maps; the gene candidates were not used in the experimental studies

| Target Gene* | Design Algorithm | Target Compound | Carbon Source | Intervention |
|---|---|---|---|---|
| YDL080C | OptKnock & OptGene | GGPP | Glucose & Galactose | Knock-out |
| YOL059W | OptKnock | GGPP | Glucose | Knock-out |
| YMR303C | OptKnock & OptGene | Acetyl-CoA | Glucose & Galactose | Knock-out |
| YDR400W | OptKnock & OptGene | Acetyl-CoA | Glucose & Galactose | Knock-out |
| YLR153C | OptKnock & OptGene | Acetyl-CoA & GGPP | Glucose & Galactose | Knock-out |
| YHR100C | OptKnock | GGPP | Glucose | Knock-out |
| YKR089C | OptKnock | GGPP | Glucose & Galactose | Knock-out |
| YML042W | OptKnock & OptForce | Acetyl-CoA | Glucose & Galactose | Knock-out |
| YDL078C | OptKnock | Acetyl-CoA | Galactose | Knock-out |
| YKL212W | OptKnock | Acetyl-CoA | Galactose | Knock-out |
| YIL006W | OptKnock & OptGene | Acetyl-CoA | Galactose | Knock-out |
| YJL097W | OptKnock | Acetyl-CoA | Galactose | Knock-out |
| YFR044C | OptGene | GGPP | Glucose | Knock-out |
| YPL268W | OptGene | Acetyl-CoA | Glucose & Galactose | Knock-out |
| YDR173C | OptGene | GGPP | Glucose & Galactose | Knock-out |
| YDR196C | OptGene | GGPP | Glucose | Knock-out |
| YGR209C | OptGene | Acetyl-CoA | Glucose & Galactose | Knock-out |
| YER086W | OptGene | GGPP | Glucose | Knock-out |
| YGR015C | OptGene & OptForce | GGPP | Glucose & Galactose | Knock-out |
| YJL200C | OptGene | GGPP | Glucose | Knock-out |
| YOR126C | OptGene | Acetyl-CoA | Glucose | Knock-out |
| YKL215C | OptGene | Acetyl-CoA | Glucose | Knock-out |
| YPL206C | OptGene | Acetyl-CoA | Glucose & Galactose | Knock-out |
| YJL005W | OptGene | Acetyl-CoA | Glucose & Galactose | Knock-out |
| YLL041C | OptGene & OptForce | GGPP | Glucose & Galactose | Knock-out |
| YER010C | OptGene | GGPP | Glucose & Galactose | Knock-out |
| YLL041C | OptGene & OptForce | Acetyl-CoA | Glucose & Galactose | Knock-out |
| YNL065W | OptGene | Acetyl-CoA | Galactose | Knock-out |
| YLR020C | OptGene | GGPP | Galactose | Knock-out |
| YLR245C | OptGene | GGPP | Galactose | Knock-out |
| YPL028W | OptGene & OptForce | Acetyl-CoA | Galactose | Knock-out |
| YPL092W | OptGene | GGPP | Galactose | Knock-out |
| YDR001C | OptGene | Acetyl-CoA | Galactose | Knock-out |
| YIL099W | OptGene | Acetyl-CoA | Galactose | Knock-out |
| YOR317W | OptForce | Acetyl-CoA | Glucose | Knock-out |
| YML059C | OptForce | Acetyl-CoA | Glucose & Galactose | Knock-out |
| YMR241W | OptForce | Acetyl-CoA | Glucose | Knock-out |
| YOR317W | OptForce | Acetyl-CoA | Glucose | Knock-out |
| YOR142W | OptForce | Acetyl-CoA | Galactose | Knock-out |
| YML120C | OptForce | Acetyl-CoA | Galactose | Knock-out |
| YOR245C | OptForce | Acetyl-CoA | Galactose | Knock-out |
| YHR063C | OptForce | Acetyl-CoA | Glucose & Galactose | Overexpression |

| | | | | |
|---|---|---|---|---|
| YGR088W | OptForce | Acetyl-CoA | Glucose & Galactose | Overexpression |
| YOR184W | OptForce | Acetyl-CoA | Glucose & Galactose | Overexpression |
| YJR024C | OptForce | Acetyl-CoA | Glucose & Galactose | Overexpression |
| YMR220W** | OptForce | GGPP | Glucose & Galactose | Overexpression |
| YNR043W** | OptForce | GGPP | Glucose & Galactose | Overexpression |
| YJL167W** | OptForce | GGPP | Glucose & Galactose | Overexpression |
| YPL069C** | OptForce | GGPP | Glucose & Galactose | Overexpression |
| YPL117C** | OptForce | GGPP | Glucose & Galactose | Overexpression |
| YMR220W** | OptForce | GGPP | Glucose & Galactose | Overexpression |

* The gene candidates suggested by OptGene or the genes of the related reactions targeted by OptKnock and/or OptForce
** These gene were not used in this study as they were previously integrated into our engineered yeast strain LRS6 (Walls et al., 2020).

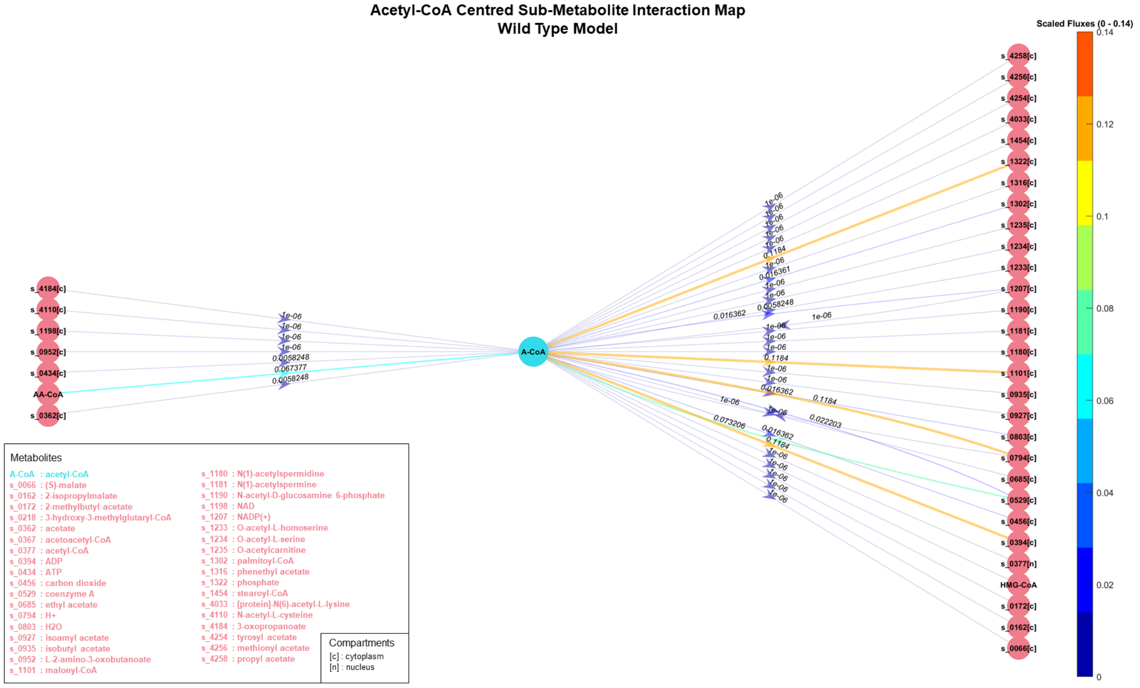

**Figure S1:** Acetyl-CoA-centred sub-metabolite interaction map of the wild-type model

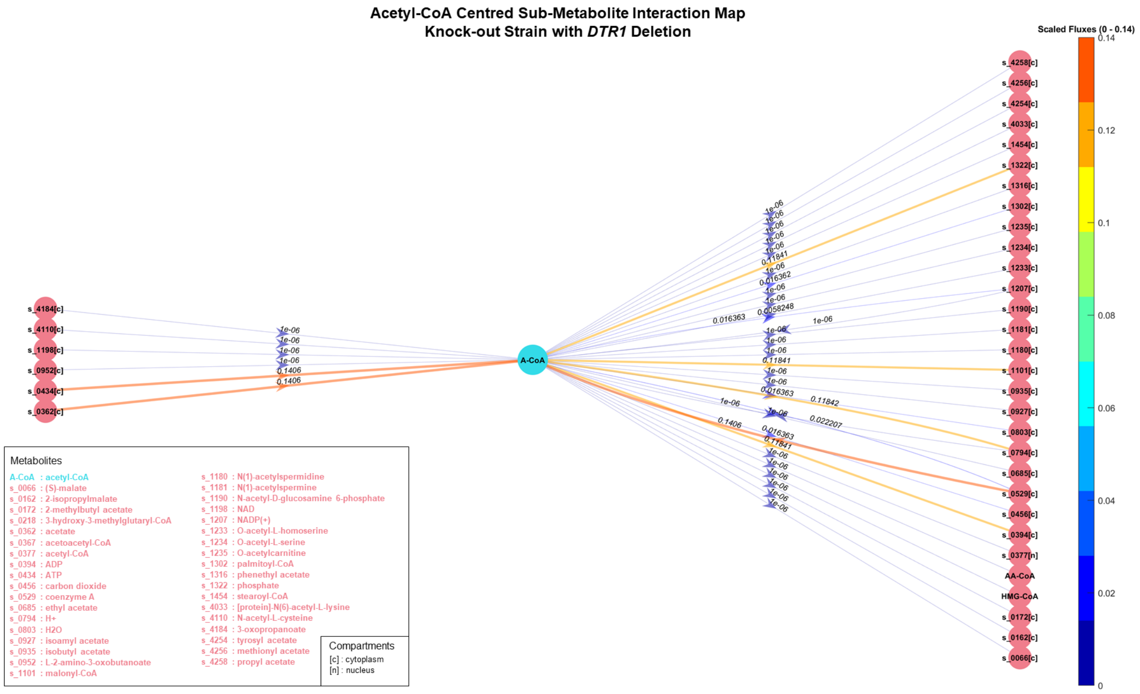


**Figure S2:** Acetyl-CoA-centred sub-metabolite interaction map of *DTR1* deleted model

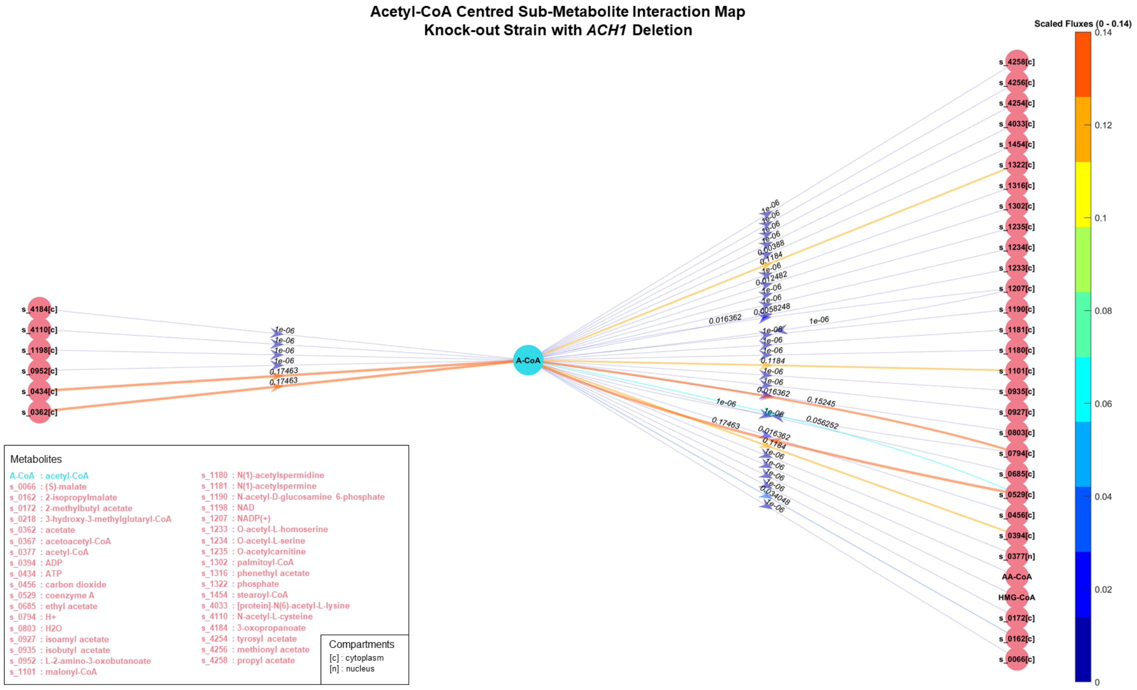


**Figure S3:** Acetyl-CoA-centred sub-metabolite interaction map of *ACH1* deleted model

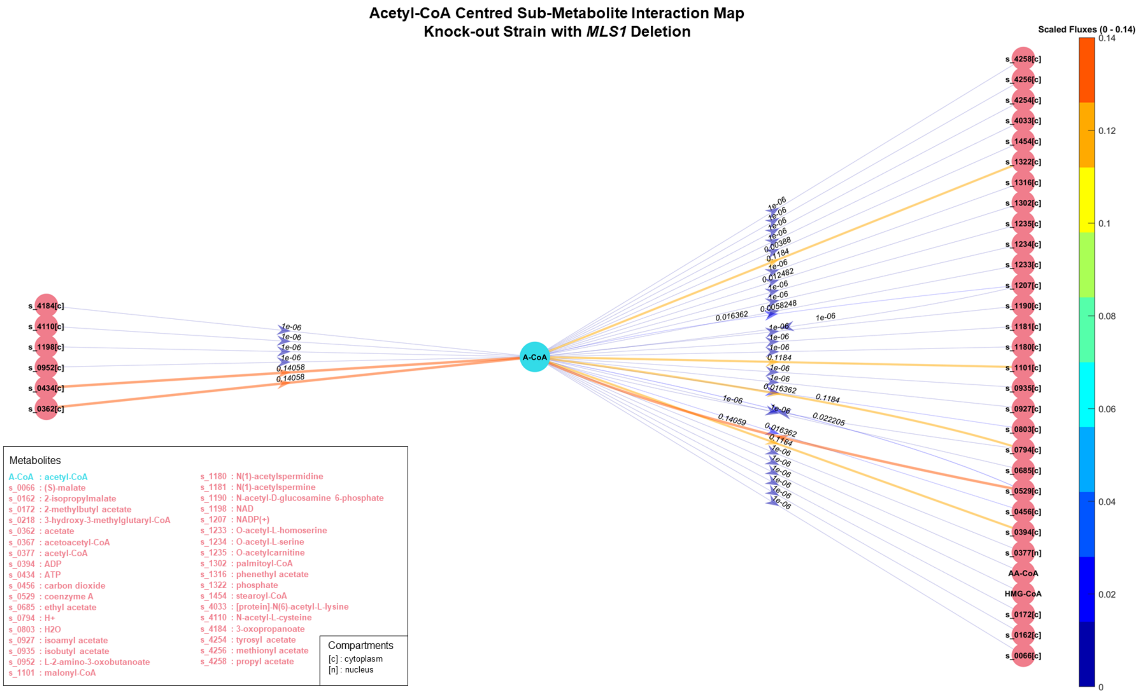


**Figure S4:** Acetyl-CoA-centred sub-metabolite interaction map of *MLS1* deleted model

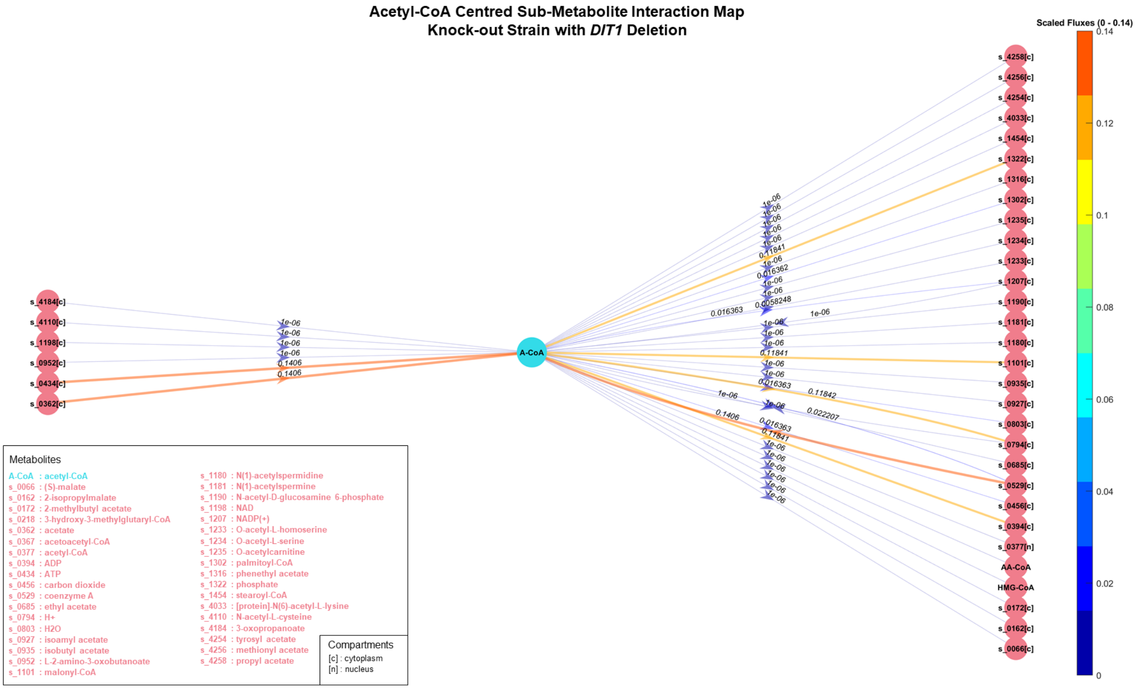


**Figure S5:** Acetyl-CoA-centred sub-metabolite interaction map of *DIT1* deleted model

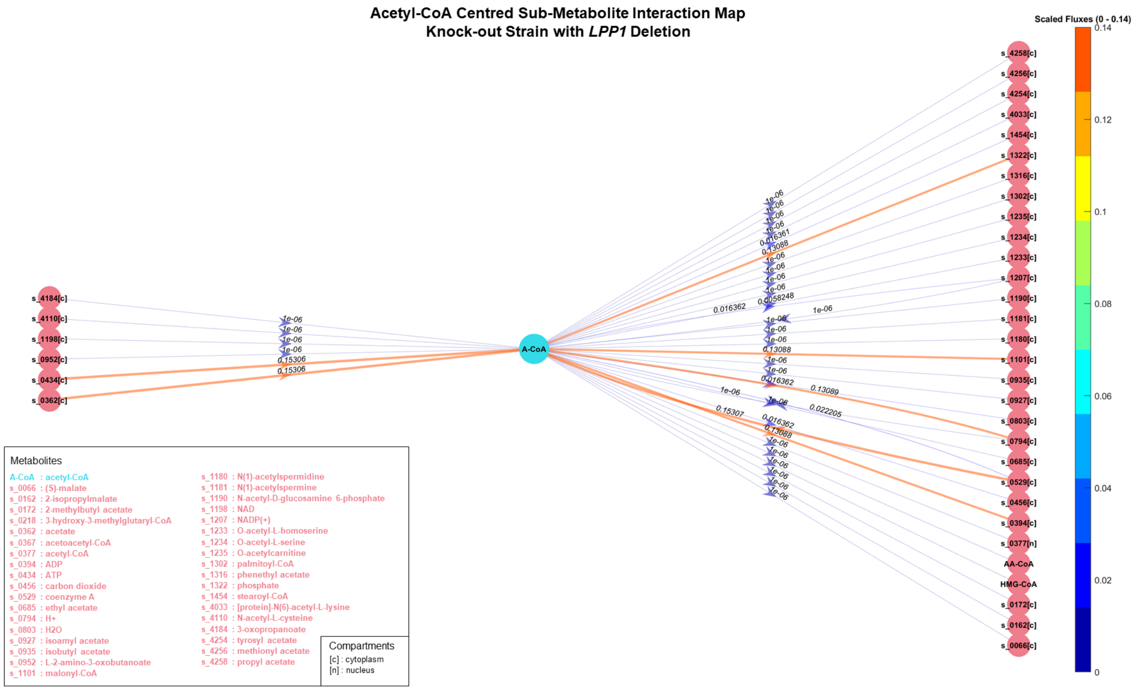


**Figure S6:** Acetyl-CoA-centred sub-metabolite interaction map of *LPP1* deleted model

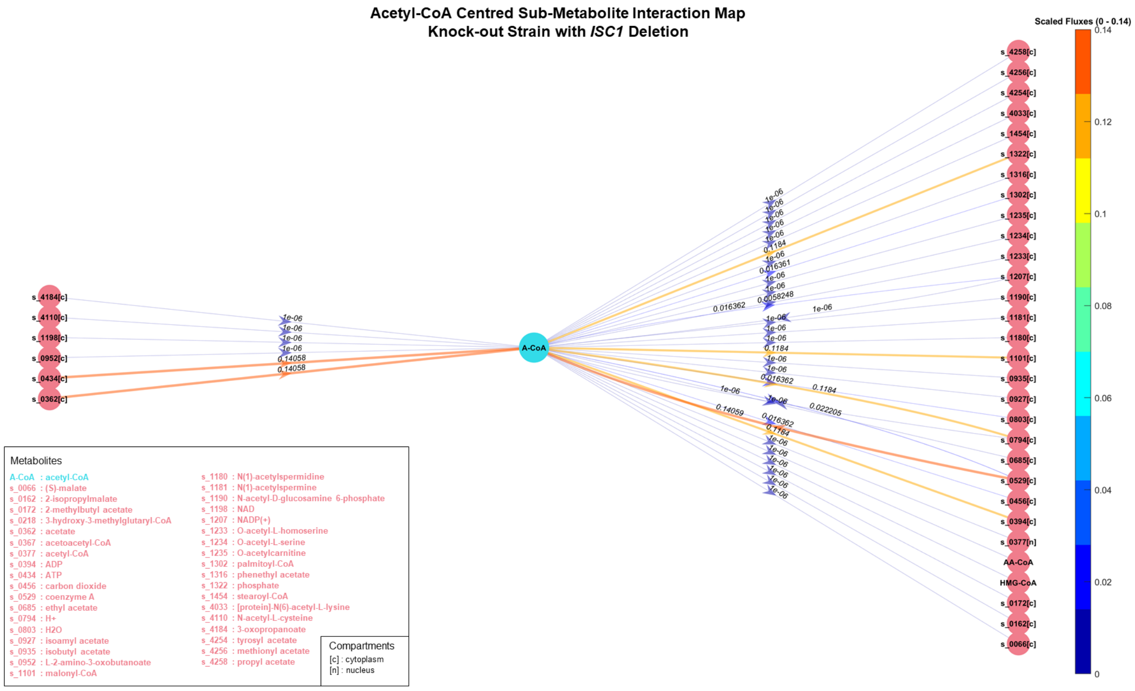


**Figure S7:** Acetyl-CoA-centred sub-metabolite interaction map of *ISC1* deleted model

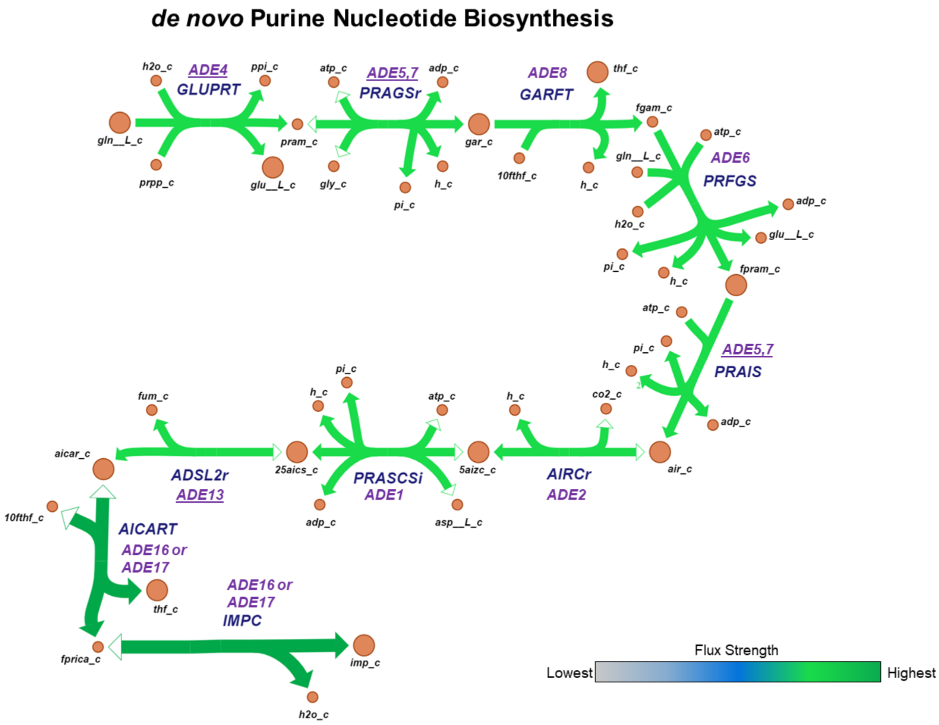


**Metabolites**

10fthf: 10-Formyltetrahydrofolate
25aics: (S)-2-[5-Amino-1-(5-phospho-D-ribosyl)imidazole-4-carboxamido]succinate
5aizc: 5-amino-1-(5-phospho-D-ribosyl)imidazole-4-carboxylate
adp: ADP
aicar: 5-Amino-1-(5-Phospho-D-ribosyl)imidazole-4-carboxamide
air: 5-amino-1-(5-phospho-D-ribosyl)imidazole
asp_L: L-Aspartate
atp: ATP
fgam: N2-Formyl-N1-(5-phospho-D-ribosyl)glycinamide
fpram: 2-(Formamido)-N1-(5-phospho-D-ribosyl)acetamidine
fprica: 5-Formamido-1-(5-phospho-D-ribosyl)imidazole-4-carboxamide
fum: Fumarate
gar: N1-(5-Phospho-D-ribosyl)glycinamide
gln__L: L-Glutamine
gln_L: L-Glutamine
glu_L: L-Glutamate
gly: Glycine
imp: IMP (C10H11N4O8P)
pi: Phosphate
ppi: Diphosphate
pram: 5-Phospho-beta-D-ribosylamine
prpp: 5-Phospho-alpha-D-ribose 1-diphosphate
thf: 5,6,7,8-Tetrahydrofolate
thf: 5,6,7,8-Tetrahydrofolate

**Reactions**

ADSL2r: Adenylosuccinate lyase
AICART: Phosphoribosylaminoimidazolecarboxamide formyltransferase
AIRCr: Phosphoribosylaminoimidazole carboxylase
GARFT: Phosphoribosylglycinamide formyltransferase
GLUPRT: Glutamine phosphoribosyldiphosphate amidotransferase
IMPC: IMP cyclohydrolase
PRAGSr: Phosphoribosylglycinamide synthase
PRAIS: Phosphoribosylaminoimidazole synthase
PRASCSi: Phosphoribosylaminoimidazolesuccinocarboxamide synthase
PRFGS: Phosphoribosylformylglycinamidine synthase

**Compartments**

_c: cytosol

**Figure S8:** *de novo* purine nucleotide biosynthesis pathway. The orange circles represent the metabolites involved in the reactions, and the genes are shown in purple. The underlined genes show the overexpressed genes. The arrows indicate the direction of the fluxes.

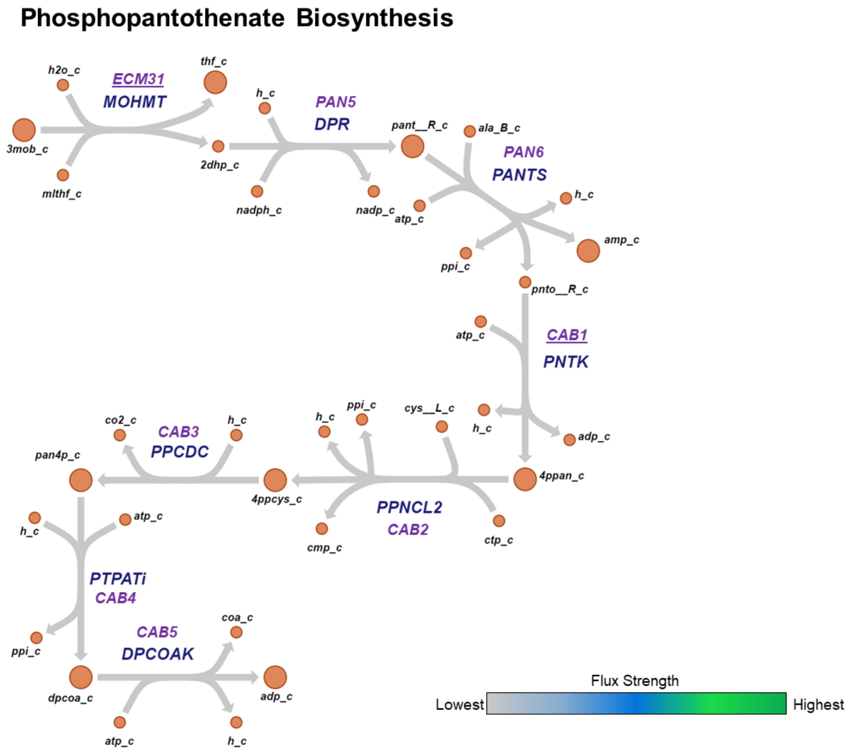


**Metabolites**

2dhp: 2-Dehydropantoate
3mob: 3-Methyl-2-oxobutanoate
4ppan: D-4'-Phosphopantothenate
4ppcys: N-((R)-4-Phosphopantothenoyl)-L-cysteine
ala_B: Beta-Alanine
amp: AMP
atp: ATP
cmp: CMP
coa: Coenzyme A
ctp: CTP
cys__L: L-Cysteine
dpcoa: Dephospho-CoA
mlthf: 5,10-Methylenetetrahydrofolate
nadp: Nicotinamide adenine dinucleotide phosphate
nadph: Nicotinamide adenine dinucleotide phosphate - reduced
pan4p: Pantetheine 4'-phosphate
pant_R: (R)-Pantoate
pnto_R: (R)-Pantothenate
ppi: Diphosphate
thf: 5,6,7,8-Tetrahydrofolate

**Reactions**

DPCOAK: Dephospho-CoA kinase
DPR: 2-dehydropantoate 2-reductase
MOHMT: 3-methyl-2-oxobutanoate hydroxymethyltransferase
PANTS: Pantothenate synthase
PNTK: Pantothenate kinase
PPCDC: Phosphopantothenoylcysteine decarboxylase
PPNCL: Phosphopantothenate-cysteine ligase
PTPATi: Pantetheine-phosphate adenylyltransferase

**Compartments**

_c: cytosol

**Figure S9:** Phosphopantothenate biosynthesis pathway. The orange circles represent the metabolites involved in the reactions, and the genes are shown in purple. The underlined genes show the overexpressed genes. The arrows indicate the direction of the fluxes.

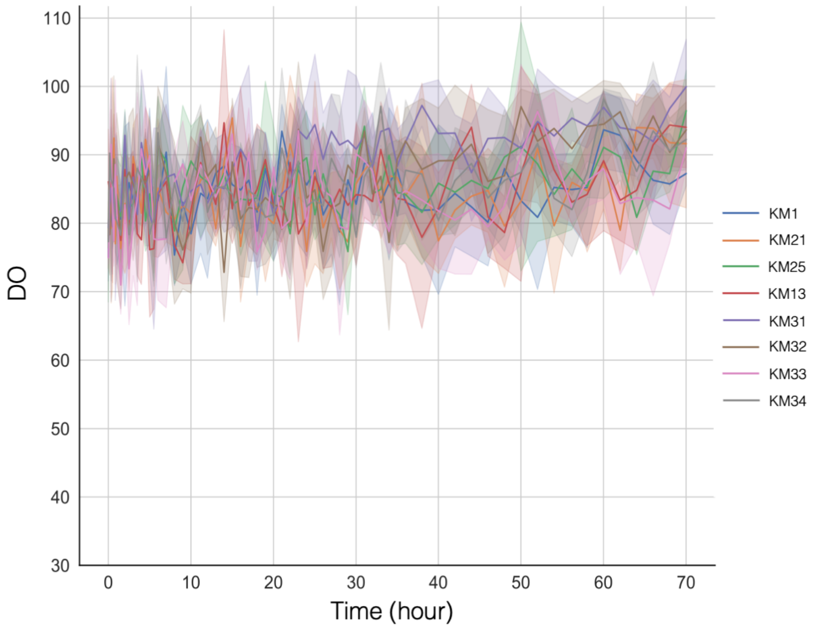


**Figure S10:** Dissolved oxygen (DO) concentrations of three-day cultures of KM1-derived strains measured by the BioLector microbioreactor system in the galactose-containing CSM

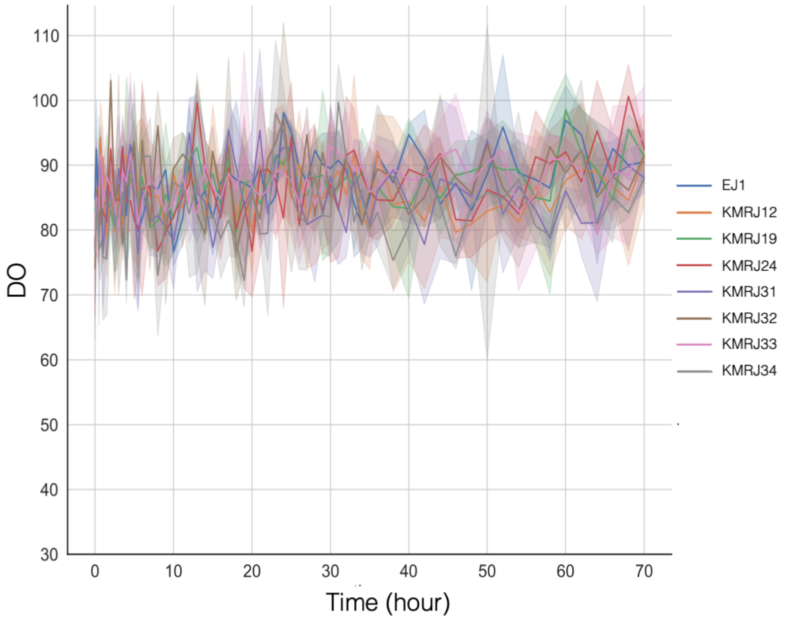


**Figure S11:** Dissolved oxygen (DO) concentrations of three-day cultures of EJ1-derived strains measured by the BioLector microbioreactor system in the glucose-containing CSM

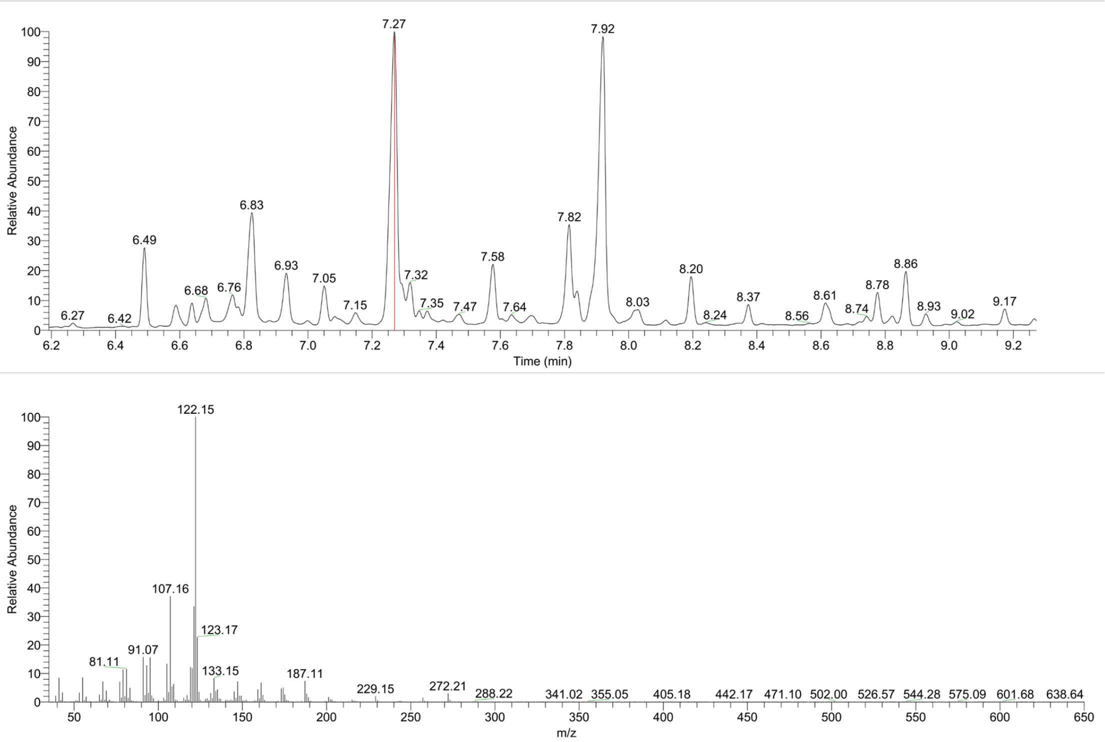


**Figure S12:** Gas chromatography shows the compounds' peaks produced by KM32 and the mass spectrum of taxadiene. The retention time of the taxadiene peak was at 7.27$^{th}$ minutes.

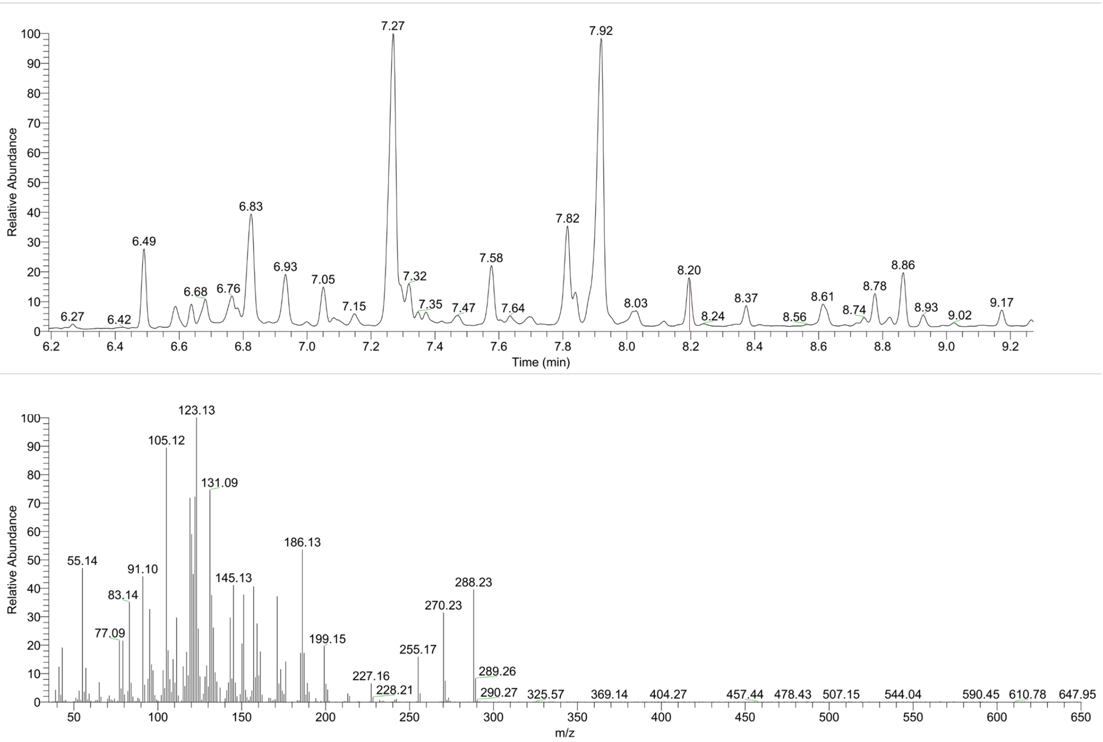


**Figure S13:** Gas chromatography shows the compounds' peaks produced by KM32 and the mass spectrum of T5α-ol. The retention time of the T5α-ol peak was at 8.20$^{th}$ minutes.

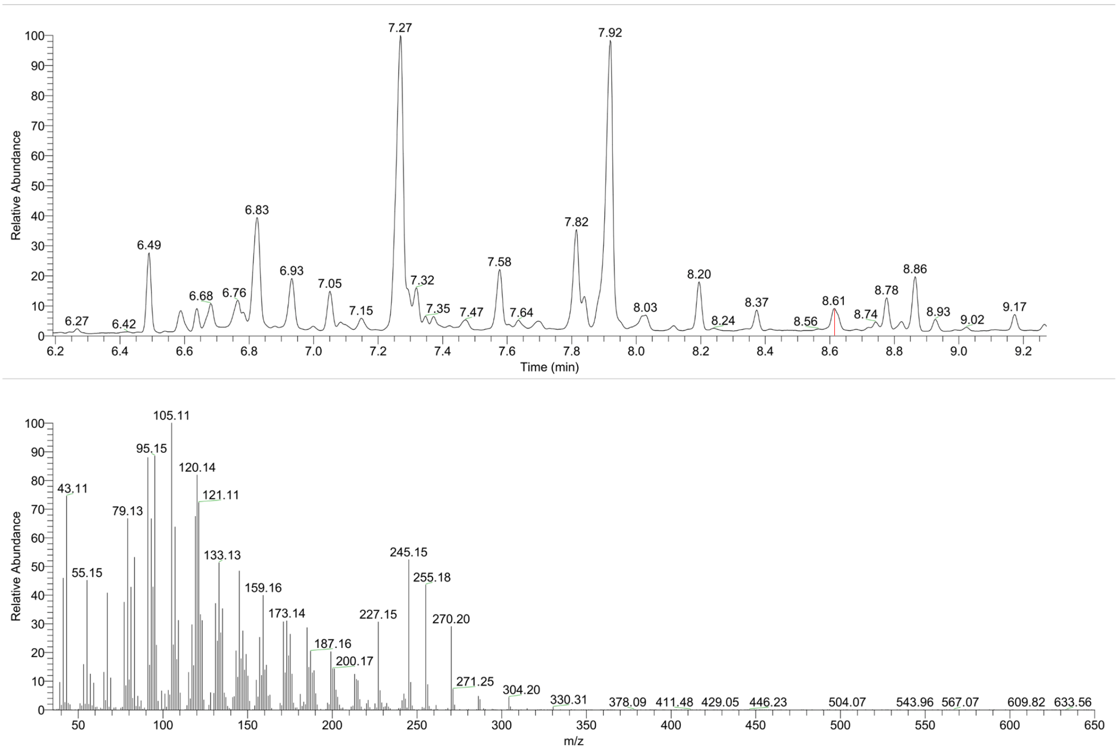


**Figure S14:** Gas chromatography shows the compounds' peaks produced by KM32 and the mass spectrum of T5αAc. The retention time of the T5αAc peak was at 8.61$^{st}$ minutes. Note that T5αAc might coelute with taxadiendiol, therefore, their peaks might be overlapped.
